# Climatic niche conservatism shapes the ecological assembly of Hawaiian arthropod communities

**DOI:** 10.1101/2021.06.22.449388

**Authors:** Jun Ying Lim, Jairo Patiño, Suzuki Noriyuki, Luis Cayetano Simmari, Rosemary G. Gillespie, Henrik Krehenwinkel

## Abstract

Spatial variation in climatic conditions along elevation gradients provides an important backdrop by which communities assemble and diversify. Lowland habitats tend to be connected through time, whereas highlands can be continuously or periodically isolated, conditions that have been hypothesized to promote high levels of species endemism. This tendency is expected to be accentuated among taxa that show niche conservatism within a given climatic envelope. While species distribution modeling approaches have allowed extensive exploration of niche conservatism among target taxa, a broad understanding of the phenomenon requires sampling of entire communities. Species-rich groups such as arthropods are ideal case studies for understanding ecological and biodiversity dynamics along elevational gradients given their important functional role in many ecosystems, but community-level studies have been limited due to their tremendous diversity. Here, we develop a novel semi-quantitative metabarcoding approach that combines specimen counts and size-sorting to characterize arthropod community-level diversity patterns along two elevational gradients across two volcanoes on the island of Hawai‘i. We find that arthropod communities between the two transects become increasingly distinct compositionally at higher elevations. Resistance surface approaches suggest that climatic differences between sampling localities are an important driver in shaping beta-diversity patterns, though the relative importance of climate varies across taxonomic groups. Nevertheless, the climatic niche position of OTUs between transects was highly correlated, suggesting that climatic filters shape the colonization between adjacent volcanoes. Taken together, our results highlight climatic niche conservatism as an important factor shaping ecological assembly along elevational gradients and suggest topographic complexity as an important driver of diversification.

## 1 Introduction

Community assembly is conceptualized as the process by which local communities assemble through the colonization and interaction of species from a larger, regional pool of species (Webb *et al.*, 2002; Cavender-Bares *et al.*, 2009; Vellend, 2016). Within this regional pool, the composition of species may be shaped by a combination of deterministic processes, such as competition or environmental filtering, or stochastic processes such as ecological drift (Hubbell, 2001; Vellend, 2016). However, while the idea that local communities are nested within the biodiversity of a larger landscape has been a hugely influential concept for understanding species composition at local scales, much less attention has been focused on how evolutionary processes shape the diversity of local communities and regional pools (Ricklefs, 2004; Rominger *et al.*, 2016).

Elevational gradients provide valuable opportunities to study how community assembly may be shaped by both ecological and evolutionary processes. Mountains have long been recognized as important biophysical arenas that are associated with high rates of diversification and endemism (Rahbek *et al.*, 2019b; Merckx *et al.*, 2015; Vasconcelos *et al.*, 2020). Climatic conditions on mountains can vary dramatically with elevation, such that a broad range of ecological conditions may occur over a relatively short distance. Given this environmental context, community assembly trajectories along elevation gradients are thus the product of complex dynamic process of environmental filtering and sorting, and the relative ability of various lineages to adapt to new climatic conditions (De Meester *et al.*, 2016). Specifically, if the climatic niches of taxa are generally conserved, then the colonization of habitats along a elevation gradient will be restricted to taxa that are already pre-adapted to prevailing environmental conditions (i.e., environmental filtering). However, if taxa are evolutionarily labile with respect to their climatic niches or show a broad climatic tolerance, then one may expect colonization of new habitats to be associated with changes in climatic niches in lineages. For instance, the colonization of mountain tops by lowland taxa, may be associated with ecological release (Yoder *et al.*, 2010). Specifically, the relaxation of selective pressures in novel environments may cause colonist lineages to expand the range of environmental conditions where they may occur (i.e., niche lability). Nevertheless, whereas both niche conservatism (Wiens, 2004; Wiens & Graham, 2005) and niche lability (Kozak & Wiens, 2010) have been argued to be potential factors that promote speciation, their relative roles in the assembly of ecological communities remains relatively unexplored.

In addition, the relative differences in isolation of different climates along elevational gradients also provides an important context for differences in community assembly trajectories and diversification along elevational gradients (Janzen, 1967; Mayr, 1947; Steinbauer *et al.*, 2016). Climates associated with higher altitudes are often insular and separated by lowland areas, whereas low-elevation areas are more well-connected by continuous habitats. Thus, the physical and environmental isolation between high-elevation communities has been hypothesized to act as an efficient barrier to dispersal and gene flow, thereby promoting speciation (Mayr, 1947; Steinbauer *et al.*, 2016; Rahbek *et al.*, 2019a, b; Salces-Castellano *et al.*, 2020). Consistent with this hypothesis, the degree of species endemism has been documented to positively correlate with elevation for many well-studied taxonomic groups, including plants (Price, 2004; Irl *et al.*, 2015), birds (Kessler & Kluge, 2008), and mammals (Heaney, 2001; Camacho-Sanchez *et al.*, 2019). In contrast, community structure is still poorly understood for groups such as arthropods (Macedo *et al.*, 2018), despite the fact that they comprise the most diverse animal group, inhabit all global ecosystems (Giribet & Edgecombe, 2012), and play pivotal roles in ecosystem function (e.g., pollination, herbivory) (Vanbergen & the Insect Pollinators Initiative, 2013). Moreover, many arthropods are sensitive to climatic variables (e.g., Halsch *et al.* 2021), making them an ideal target group to study the role of elevation on community and diversity patterns.

To understand the role of niche conservatism and elevation-driven isolation on the assembly of communities, we characterize arthropod communities of native *Metrosideros* forests along two elevational gradients across two different volcanoes on the oceanic island of Hawai‘i, using a novel semi-quantitative metabarcoding approach, based on size sorting and counting of all sampled individuals. The island of Hawai‘i provides an ideal landscape by which the process of community assembly across elevational gradients may be studied for two main reasons: firstly, active volcanism and frequent lava flows on the island would have produced blank slates by which biological processes may then act upon, thus proving us an opportunity to study the process of community assembly *de novo*(Carson *et al.*, 1990; Rominger *et al.*, 2016); secondly, the island is composed of multiple volcanic shields of different geological ages, and so populations on older volcanoes act as sources for geologically younger substrates, thus controlling for initial source pools (Shaw & Gillespie, 2016).

We hypothesize that 1) populations across the two transects will have similar climatic preferences due to niche conservatism. As higher elevation areas are more geographically and climatically isolated than lowlands, we also hypothesize that 2) beta-diversity would increase with higher elevation. Lastly, we predict that 3) the greater isolation of higher elevations would lead to higher genetic differentiation between populations at higher elevations compared to populations at lower elevations.

## 2 Methods

### 2.1 Transects, sampling sites and sample collection

Arthropod communities were sampled along two elevational transects on the Big Island of Hawai‘i. The first transect was on the north-eastern slope of Mauna Kea (hereafter the Laupāhoehoe transect). The second transect was on the eastern slope of Mauna Loa (hereafter the Stainback transect). Both transects spanned approximately 1000 m of elevation, but the putative ages of the substrates at both sites are substantially different given the volcanic shields upon which they are situated. The Laupāhoehoe transect traversed relatively ancient lava flows between 5,000 to 11,000 years old, whereas the Stainback transect cut across a mosaic of substrates of largely younger age, the large majority ranging between 200 – 1,500 years old (Sherrod *et al.*, 2007). Mauna Kea is considerably older than Mauna Loa and is the most likely source pool for communities on Mauna Loa due to its proximity.

The canopy at all sampling sites was largely dominated by the endemic tree, ‘ō‘hia lehua (*Metrosideros polymorpha*), while the understory was dominated by the hāpu‘u tree ferns, *Cibotium glaucum*. The presence of invasive plants in the undergrowth was minimal and disturbed areas were avoided. Arthropod communities were thus expected to be dominated by native taxa. We started sampling at the lowest elevation of native Metrosideros forest, at around 800 m. Moving up about 1000 m from this elevation, we sampled arthropods at a total of 23 sites along the Laupāhoehoe transect and 36 sites along the Stainback transect. Replicated sites at different elevations within each transect were sampled, making a total of 59 plots. Both transects showed a congruent linear decline in mean annual temperature with increasing elevation (Supplementary Fig. S1). Total annual precipitation decreased linearly with elevation along the Laupāhoehoe transect, but the Stainback transect is much wetter at lower elevations and displays a much stronger mid-elevational precipitation gradient (Supplementary Fig. S1). GPS coordinates of sampling site centroids were logged and sampling took place in a 15 m radius around the centroid. Arthropods were collected by beating native undergrowth vegetation for a total of four person-hours per site. Undergrowth included shrubs as well as small trees, including young Metrosideros trees, at all sites. The canopy of mature trees was not sampled due to their height. The vegetation was carefully beat with a 1 m PVC pipe and falling arthropod specimens collected on white beat sheets of 1 m^2^ size. Specimens were then collected from the sheets using aspirators. Specimens were subsequently stored in ethanol and further processed at the University of California Berkeley.

### 2.2 Specimen sorting, DNA extraction and library preparation

Specimens of large body size will contribute disproportionately more DNA than small ones, qualitatively and quantitatively biasing the DNA sample (Vandeputte *et al.*, 2017). To ameliorate this bias, we first sorted samples into body size categories. Specimens from all collection sites were measured using graph paper under binocular microscopes and separated into four size categories and counted: 0 – 2 mm, 2 – 4 mm, 4 – 7 mm and > 7 mm. In many sites, Collembola were highly abundant, and thus potentially make up a considerable proportion of the collected biomass. Collembola were thus also separated from the rest of the taxa and counted so their contribution to the final read abundance may be accounted for.

All size sorted specimens were transferred into 96 well 2 ml block plates and DNA extraction performed using the Qiagen Puregen Kit (Qiagen, Hilden, Germany) with the following modifications. A single 5 mm stainless steel bead and 1 ml lysis buffer and 5 *μ*l Proteinase K (Qiagen, Hilden, Germany) were added to each sample. The block plates were then sealed with reusable silicone mats and all specimens ground finely on a 2010 Geno/Grinder (SPEX Sample Prep, Metuchen, USA) at 1,300 hz for 2 minutes. Protein precipitation was then performed and the supernatant transferred to a new plate. DNA was then isolated from the samples on a Pipetting robot (Beckman Coulter, Indianapolis, USA) by magnetic bead enrichment (Bioneer, Kew East VIC, Australia). The final cleaned DNA sample was eluted into 50 *μ*l TE Buffer. The bead-based DNA isolation removes PCR inhibitors efficiently, yielding DNA, which can be directly used for PCR.

To minimize PCR amplification bias, we used universal and highly degenerate Primers targeting a short stretch of the mitochondrial Cytochrome Oxidase Subunit I (COI). We used two primer pairs targeting an overlapping COI region, to additionally mitigate bias (Leray *et al.*, 2013; Gibson *et al.*, 2014). We have previously tested these primers extensively with Hawaiian arthropod communities and, in combination, they reliably amplify the majority of tested taxa (de Kerdrel *et al.*, 2020). Libraries were prepared using a dual indexing strategy as described in Lange *et al.* (2014). The libraries were then quantified on a 1.5% agarose gel and pooled in approximately equal amounts based on gel intensity. The pooled sample was then cleaned up from residual primer by 1 X Ampure Beads XP (Beckman Coulter, Indianapolis, USA). The cleaned libraries were sequenced on an Illumina MiSeq using V3 chemistry and 600 cycles at the Center for Comparative Genomics of the California Academy of Sciences. Negative control PCRs were amplified and sequenced along with the regular samples.

### 2.3 Sequence analysis

Demultiplexed reads were merged using PEAR (Zhang *et al.*, 2014), with a minimum quality of 20 and a minimum overlap of 50 bp. The merged reads were then additionally quality filtered for sequences with a minimum of 90% bases with a quality of 30 and then transformed to fasta using Fastx-toolkit (Gordon & Hannon, 2010). PCR primer sequences were trimmed off from all merged reads using the functions grep and sed in UNIX. The two overlapping amplicons were then trimmed to the exact same regions for further analysis. All sequence data was then merged and dereplicated using USEARCH (Edgar, 2010, 2016). USEARCH was used to generate zero radius OTUs (hereafter zOTUs) using the unoise3 command, and 3% radius OTUs (hereafter OTUs) using the `cluster_otus` command. Chimeras were removed during OTU clustering.

All resulting OTU sequences were translated in MEGA (Tamura *et al.*, 2013) and only sequences with intact reading frames retained. All remaining OTU sequences were then compared to Genbank using BLASTn (Altschul *et al.*, 1990), with a maximum number of 10 target sequences. Only sequences identifying as arthropods were retained. By blasting, order level taxonomy could be assigned to the majority of OTUs. We assigned order level status if all 10 BLAST hits were assigned to the according order with > 90% similarity. The majority of OTUs could be classified this way. For the OTUs that remained unassigned, their taxonomic order was determined by their phylogenetic placement in a maximum likelihood phylogeny of all OTUs.

### 2.4 Read rarefaction and controlling for sampling bias

An OTU table was constructed in USEARCH by mapping the raw reads back to the cleaned zOTU sequence file. Separate samples were mapped for sampling site, size category and amplicon. The initial OTU table thus contained a total of 10 entries per sample site (five sorted categories for two loci). We rarefied read abundances of samples by the number of individuals that were sorted in each size category for each site. For each size category at each site, reads were randomly sub-sampled based on the number of individuals recorded for that size category at that site. The number of reads per individual we sub-sampled was chosen as the largest number of reads such that every single site - size category sample could be sub-sampled at the same level.

The rarefaction procedure ensures that the sampling sequencing effort (i.e., the total number of reads) is effectively equalized across body size classes and sites. Rarefaction was performed 100 times for each of the two amplicons separately. Rarefied read abundance for each OTU was summed for each marker to produce 100 rarefied OTU datasets. Rarefaction was performed using custom scripts based on the `rrarefy` function in the vegan R package (v 2.5-7, Oksanen *et al.* 2020). Finally, the datasets for the two markers for all body size classes were combined. The final OTU table thus contained a single entry per sampling site, effectively merging the size classes, as well as the two markers.

Using the BLASTn assignments, we identified OTUs from six arthropod orders for analysis: Araneae, Coleoptera, Hemiptera, Lepidoptera, Orthoptera, and Psocoptera. We excluded reads from OTUs that belonged to other orders as they were poorly represented (i.e., very few OTUs sampled) or from orders that may be represented in a biased manner in a beat sample (e.g., Diptera and Hymenoptera, which are able to quickly escape or fly off the beat sheet). The abundances and diversity for such groups in beat samples may thus be biased or idiosyncratic (de Kerdrel *et al.*, 2020); however, we included Lepidoptera because these are well represented by non-flying larval stages. We additionally excluded OTUs from the orders Isopoda, Collembola, Blattodea, Myriapoda and Amphipoda, because specimens sampled from those orders were of non-native species.

### 2.5 Elevation, climate and geographic distance on alpha and beta-diversity patterns

To evaluate alpha diversity patterns, we calculated OTU richness and Shannon’s index for all sites and for each taxonomic order separately. To evaluate beta-diversity patterns between transects, we calculated abundance-weighted and non-weighted (binary) Bray-Curtis dissimilarity values between sites on Laupāhoehoe and Stainback transects. We first performed a non-metric dimensional scaling of all sites on both transects using the mean read abundance of OTUs across all 100 rarefied datasets. In addition, to quantify how beta-diversity changes with elevation, we compared sites from each transect that are within 100 m elevation of each other. We tested whether dissimilarity values increase with increasing elevation (calculated as the mean elevation of the two sites compared) using Spearman’s correlation tests. We performed this analysis with presence-absence and abundance-weighted Bray-Curtis dissimilarity values, and for each taxonomic order separately.

In addition, we hypothesize that increasing isolation with higher elevations should also lead to increased genetic differentiation within OTUs. To test this, we additionally look at turnover in zero-radius OTUs (zOTU) among OTUs with increasing elevation. This is based on the assumption that dissimilarity in zOTUs for a given OTU may provide a proxy for the degree of genetic differentiation within the OTU between transects. Using the mean read abundance of zOTUs across 100 rarefied datasets, we first calculated using Bray-Curtis dissimilarity (read abundance-weighted and unweighted) in zero-radius OTUs for each OTU between sites. We then calculated the average dissimilarity across OTUs between sites from each transect within 100 m elevation of each other. Dissimilarity with elevation was then tested with Spearman’s correlation tests. All Bray-Curtis dissimilarity values (weighted and non-weighted) were calculated using the `vegdist` function of the vegan R package.

To evaluate the potential of beta-diversity patterns for predicting connectivity within transects, we used an optimization method in order to calculate resistance surfaces in the R package ResistanceGA v. 4.0 (Peterman, 2018). Specifically, based on a genetic algorithm and a maximum likelihood population effects parameterization (MLPE) model (i.e., a mixed effects model framework), this approach optimizes resistance surface values such that effective distances between sampling locations across the landscape. It has been demonstrated that MLPE mixed models performed better than Mantel’s tests and multiple regression on distance matrices, as they account for non-independent variables and are less prone to error (Row *et al.*, 2017). These effective distances were calculated as the random-walk commute time in the gdistance R package (van Etten, 2017), which is a multi-path distance measure equivalent to resistance distance (Kivimäki *et al.*, 2014), as calculated by Circuitscape (McRae *et al.*, 2008). The R package ResistanceGA is unique among available approaches for landscape resistance modelling in that it is a true optimization of landscape resistance values, avoiding subjectivity in assigning resistance values (Peterman *et al.*, 2014; Peterman, 2018). We used the all_comb function in ResistanceGA to optimize all single- and multiple-surface combinations of resistance surfaces (up to two surfaces within a model). For multiple-surface optimization, resistance surfaces were combined during optimization to create composite resistance surfaces. We used the Akaike information criterion corrected for small-sample size (AICc) to assess support for optimized resistance surfaces, and identify the best set of fitted MLPE mixed models according to the criterion of ∆AICc < 2 (Burnham & Anderson, 2002). Complementarily, we evaluated the model selection procedure for each layer and the combinations of layers by fitting each surface to a random selection of 75% of samples without replacement for 10,000 iterations in order to obtain the percentage of top ranked model occurrences (Peterman, 2018).

In terms of environmental data, we used two climatic layers, temperature and precipitation, obtained from the climatic rasters for the Hawaiian islands (Giambelluca *et al.*, 2013, 2014). Layers were analyzed as independent surfaces as well as multi-surface RasterStack objects to consider one- and two-way interactions among all climatic features. We assessed the effects of these resistance surfaces, and compared with an isolation-by-distance hypothesis, on community-level pairwise distance statistics (see Salces-Castellano *et al.* 2020). As the analysis was computationally expensive, we performed the analysis of five randomly selected rarefied OTU datasets. The ResistanceGA analyses were run at El Teide high-performance computing cluster facility (Instituto Tecnológico y de Energías Renovables).

### 2.6 Estimating the role of niche conservatism in community assembly

We quantified the climatic niche (i.e., mean annual temperature (TEMP) and total annual precipitation (PREC)) of each OTU on each transect. PCR amplification bias will lead to skewed abundance estimates, when comparing different species within a community. A single species, however, will perform relatively similarly in different PCR reactions (Amend *et al.*, 2010; Krehenwinkel *et al.*, 2017). The sample size corrected relative abundance differences within a single species, but between different sampling sites should thus be accurate. By rarefying our dataset by the number of specimens collected at each site, different sites are directly comparable. We used this to estimate the relative abundance for different OTUs across the sampled gradients. The relative abundance of a species along the transect should reflect its climatic niche "position", with optimal conditions yielding high abundances of the species. Under an assumption of “climatic equilibrium” (*sensu* Soberón 2007), to estimate the niche position for each OTU, we estimated the median climatic value of reads at each sampling locality (i.e., half of reads were recovered from sites that were either higher or lower than this value). To exclude OTUs that are found at only a few sites, we performed this analysis on OTUs that occurred in at least 5 sites per transect. We additionally excluded OTUs whose interquartile range overlapped with the upper and lower climatic bounds sampled for each site (i.e., at least 25% of all reads occurred at sampling localities that represent either the upper or lower bounds for each climatic variable for each site). This was to reduce inaccuracies in the estimation of niche positions of such OTUs as their elevational ranges may appear truncated by our sampling extent. We then tested whether the niche position of OTUs was correlated between the two sites using Spearman’s correlation tests. We repeated our analysis across the 100 rarefied datasets. Significance for each correlation test was evaluated by permutation. To evaluate the sensitivity of the results to our rarefaction procedure, we performed additional analyses where we only included OTUs that occurred in all rarefied datasets as well as a higher threshold for OTU inclusion: OTUs that occurred in at least 10 sites per transect. Values of mean annual temperature and total annual precipitation were obtained from the climate atlas of Hawai‘i (Giambelluca *et al.*, 2013, 2014). We did not perform this analysis for each order separately, as the number of OTUs that met our criteria was too low.

## 3 Results

### 3.1 Metabarcoding dataset and alpha diversity patterns

We collected a total of 39,092 specimens along the Stainbeck transect and 20,931 across the Laupāhoehoe transect. Along the Stainback transect, an average of 1,086 specimens were collected at each site, with 17% of specimens in the smallest size category of 0–2 mm, 15% in the 2–4 mm category, 6% in the 4–7 mm category, 4% in the > 7 mm categorym and the remaining in the “Collembola” category. Along the Laupāhoehoe transect, 910 specimens were collected per site on average. Of these, 27% fell into the smallest size category of 0–2 mm, 41% into the 2–4 mm category, 9% into the 4–7 mm category, and 5% into the > 7 mm category, with the remaining in the “Collembola” category.

Across all rarefied datasets, average OTU richness was higher in sites along the Laupāhoehoe transect (mean = 109) compared to sites along the Stainback transect (mean = 83) (Welch’s t-test, p < 0.05) (Fig. 1b). This was largely true for most taxonomic orders except for Araneae (Supplementary Fig. S2). OTU richness showed a hump-shaped pattern with elevation along Laupāhoehoe, with diversity peaking between 1,100 - 1,300 m a.s.l. (above sea level; Fig. 1b). On Stainback, OTU richness gradually increased with greater elevation but reached a plateau around 1,100 m elevation (Fig. 1b). Shannon diversity, however, was not significantly different between sites (Welch’s t-test, p = 0.29; Supplementary Fig. S3). Relative proportion of OTUs from various taxonomic orders was generally constant with elevation, with the exception of Orthoptera, which declined in diversity and proportional read count with increasing elevation (Supplementary Fig. 4).

**Figure 1.**
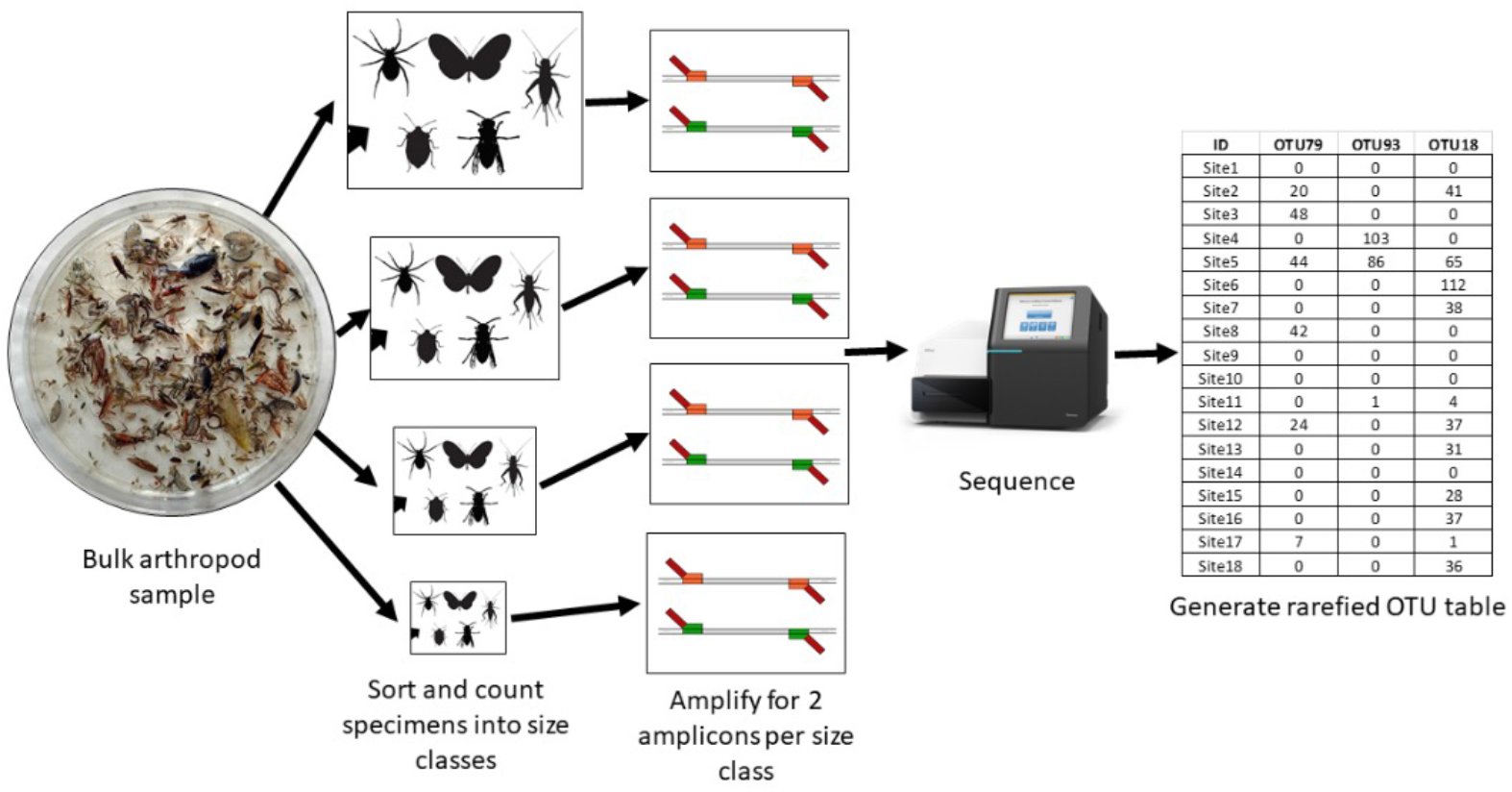
Processing pipeline to control for sampling effort and body size of specimens. Bulk samples were sorted into size categories (see Methods) and specimens counted, then amplified for two COI amplicons and sequenced. The final dataset was then rarefied based on initial specimen abundances for each size category and marker separately and then combined.

**Figure 2.**
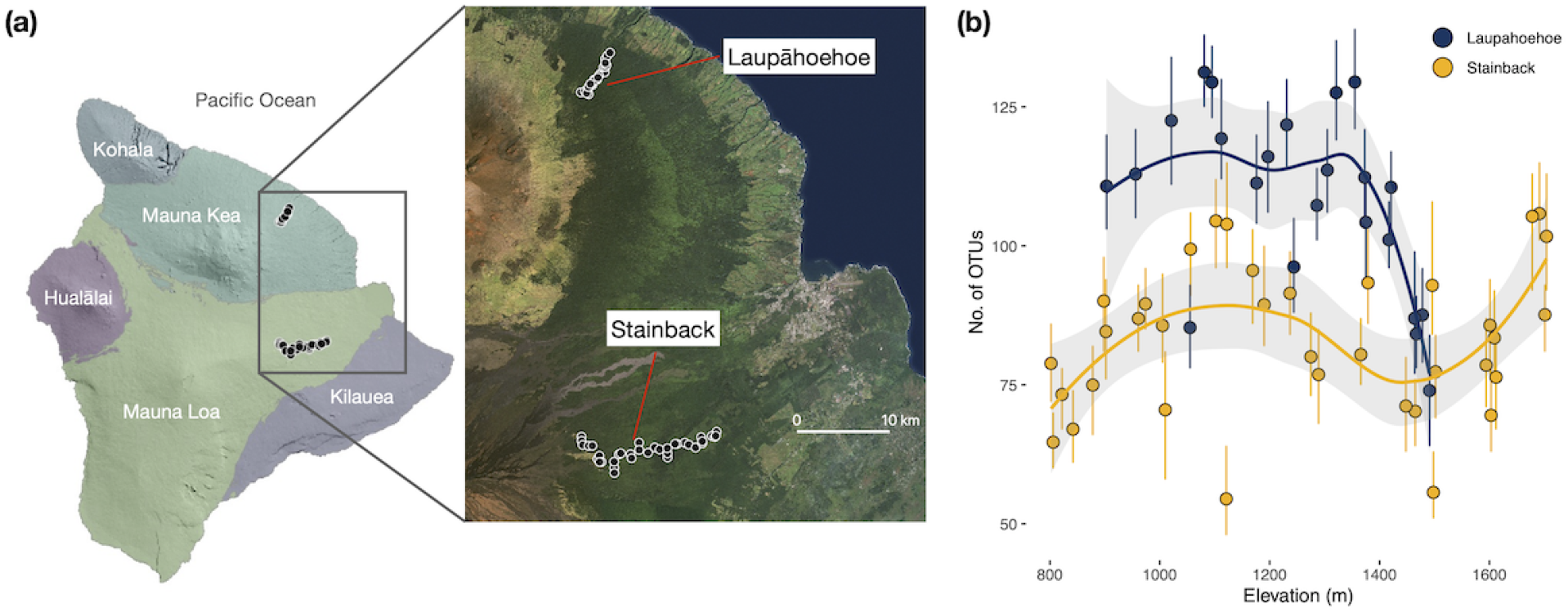
Geographic location (a) and patterns of OTU richness (b) across two elevational transects on the volcanoes Mauna Kea (Laupāhoehoe) and Mauna Loa (Stainbeck) on the island of Hawai‘i. Points represent the average OTU richness for each site across rarefied datasets, whereas error bars represent the minimum and maximum OTU richness across rarefied datasets. Lines (mean) and shaded area (95% confident intervals) depict trends in OTU richness for each transect and were generated using a loess smoothing function on average OTU richness of sites in each transect.

### 3.2 Effect of elevation, climate and geographic distance on beta-diversity patterns

Between transects, arthropod communities were compositionally more similar at lower elevations than higher elevations. The NDMS plot shows a V-shaped pattern of community differentiation along the transects suggesting that community composition between transects was more similar at low elevations and gradually diverged at higher elevations (Fig. 3a). Abundance-weighted dissimilarity of sites between transects was highly positively correlated with elevation (Spearman’s correlation, *ρ* = 0.65; p < 0.001)(Fig. 3b). The pattern of higher compositional dissimilarity between transects at higher elevations was also true when taxonomic orders were analyzed separately (Spearman’s *ρ* = 0.19 – 0.62, p < 0.05; Supplementary Fig. S6). We found a significant increase in community dissimilarity with increasing elevation for all analyzed six orders of arthropods. Unweighted estimates of community dissimilarity (i.e., presence-absence, *ρ* = 0.68, p < 0.001; Supplementary Fig S5a,b) generally show the same patterns of community differentiation as abundance-weighted measures. However, the pattern was less clear when orders were analyzed separately (Supplementary Fig. S6). Unweighted dissimilarity within Lepidoptera and Araneae assemblages respectively do not appear to be correlated with elevation between the transects (Lepidoptera *ρ* = 0.14, p = 0.08; Araneae *ρ* = −0.04, p = 0.66). For Psocoptera, the pattern even reversed, showing a decreasing turnover with elevation (*ρ* = −0.25, p < 0.05; Supplementary Fig. S5c).

**Figure 3.**
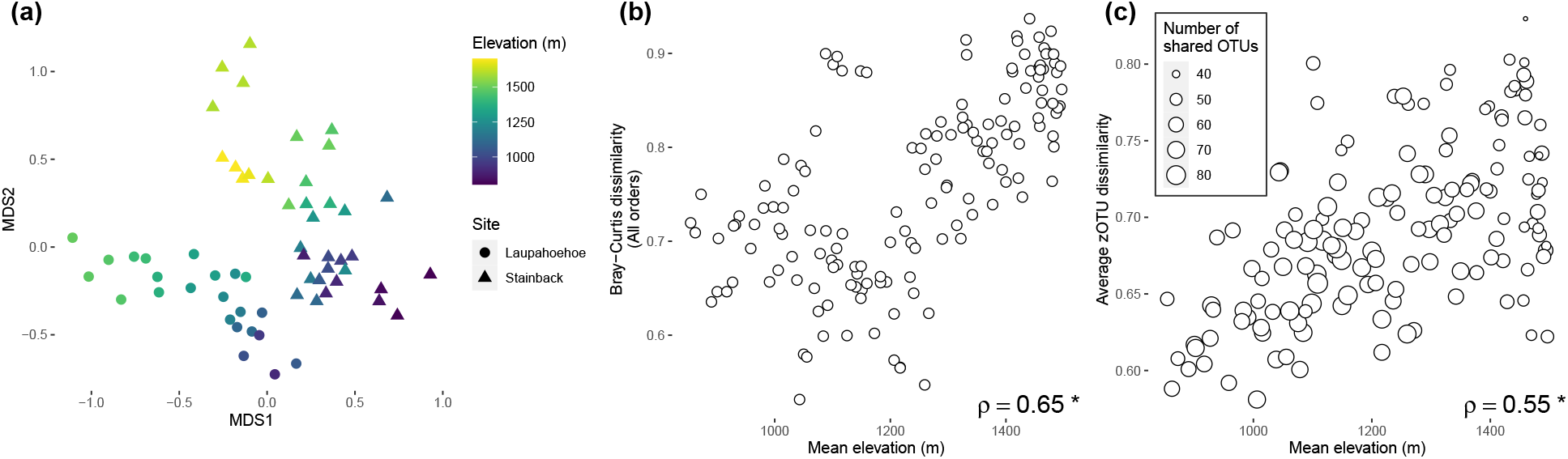
The effect of elevation on community compositional dissimilarity (unweighted Bray-Curtis dissimilarity) between sites on Laupāhoehoe vs. Stainback transects. a) Non-metric dimensional scaling analysis of sampling localities across sites (triangles = Stainback, circles = Laupāhoehoe). NMDS stress value = 0.18. b) Pairwise dissimilarity between Laupāhoehoe and Stainback sampling localities against the mean elevation of localities compared. Only sampling sites between transects that were within 100 m elevation of each other were compared. Arthropod communities are increasingly dissimilar between Laupāhoehoe and Stainback with higher elevation (Spearman’s correlation, *ρ* = 0.65, p < 0.05). c) Average unweighted dissimilarity in zOTU composition of OTUs between sites on different transects increases with increasing elevation (*ρ* = 0.55, p < 0.05).

The number of shared OTUs between transects decreased with elevation. However, average abundance-weighted dissimilarity in zOTU composition within OTUs increased with increasing elevation between sites (Fig. 3c). This pattern was stronger when considering just the presence-absence of zOTUs for each OTU (Fig. S5c).

Resistance landscape analyses of the climatic variables suggests that patterns of arthropod community dissimilarity are globally best explained as a function of climate rather than geographic isolation (Supplementary Table 1, Fig. 4). While simple geographic distance was a poor predictor of community dissimilarity patterns for most taxonomic orders along the Stainback transect, geographic isolation between localities along the elevational gradient appears to play a significant role along the Laupāhoehoe transect for only two orders: Araneae and Lepidoptera (Fig. 4).

**Figure 4.**
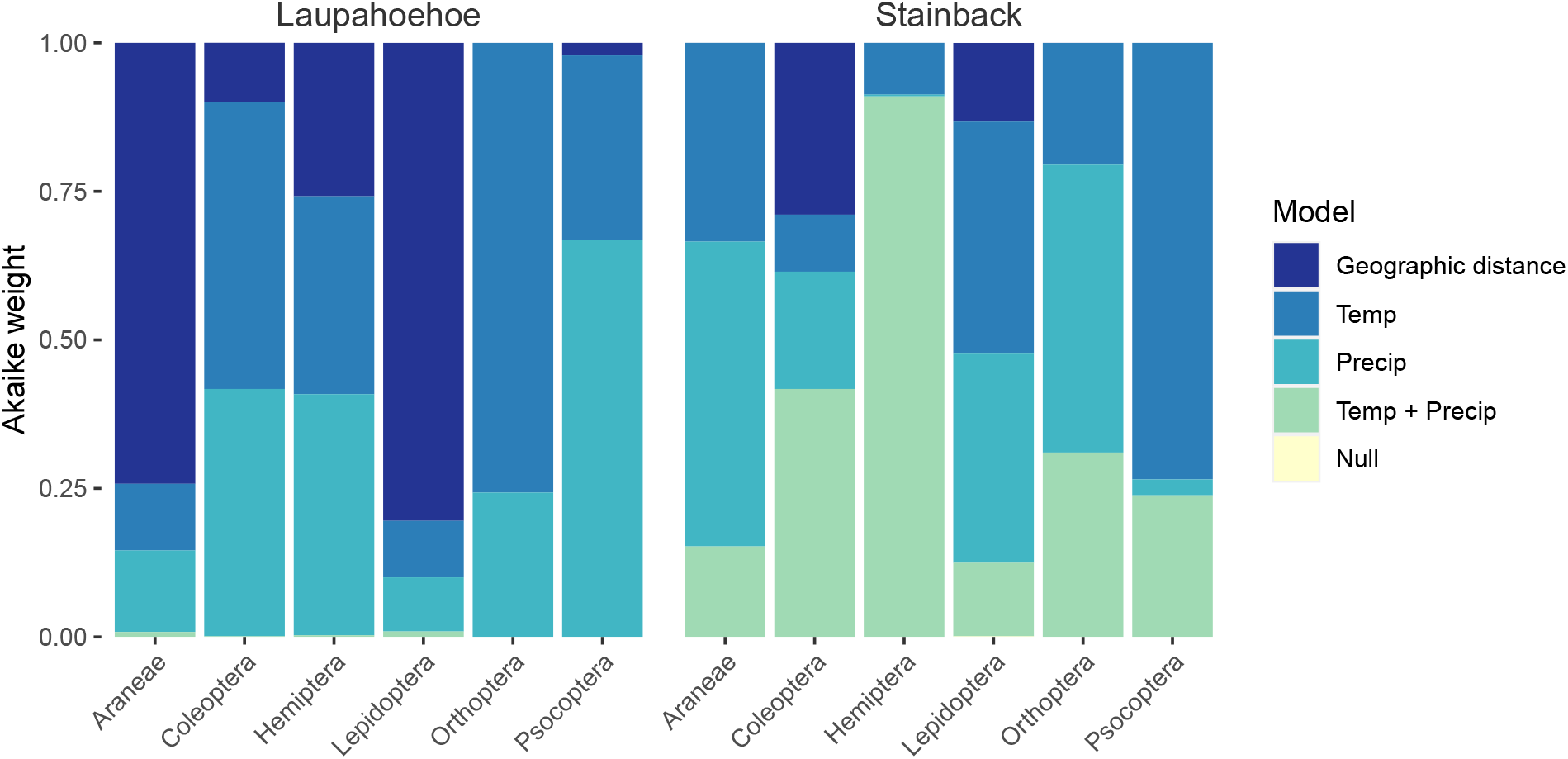
Relative explanatory power (Akaike weights) of geographic distance and differences in climate in explaining beta-diversity patterns for arthropod communities of different taxonomic orders. Akaike weights for each model are averages across models of five randomly subsampled rarefied datasets.

### 3.3 The role of climatic niche conservatism in ecological assembly

Climatic niches of OTUs were highly correlated between transects (Fig. 5). The conservation and predictability of niche space was found for both precipitation and temperature. Mean Spearman’s correlation coefficient, across rarefied datasets, for temperature was 0.39 (number of OTUs, n = 86 – 99); whereas the mean correlation for precipitation was 0.26 (n = 50 – 59). When we analyzed only OTUs that occurred in all rarefied datasets, the mean correlation coefficient across rarefied datasets for temperature and precipitation was 0.31 (n = 67) and 0.27 (n = 45), respectively. Differences in sample size between temperature and precipitation tests can be explained by differences in the precipitation gradient between the transects. Correlations in climatic niches were weaker when we limited our analysis to OTUs that were found in at least 10 sites in each transect (Supplementary Fig. S7).

**Figure 5.**
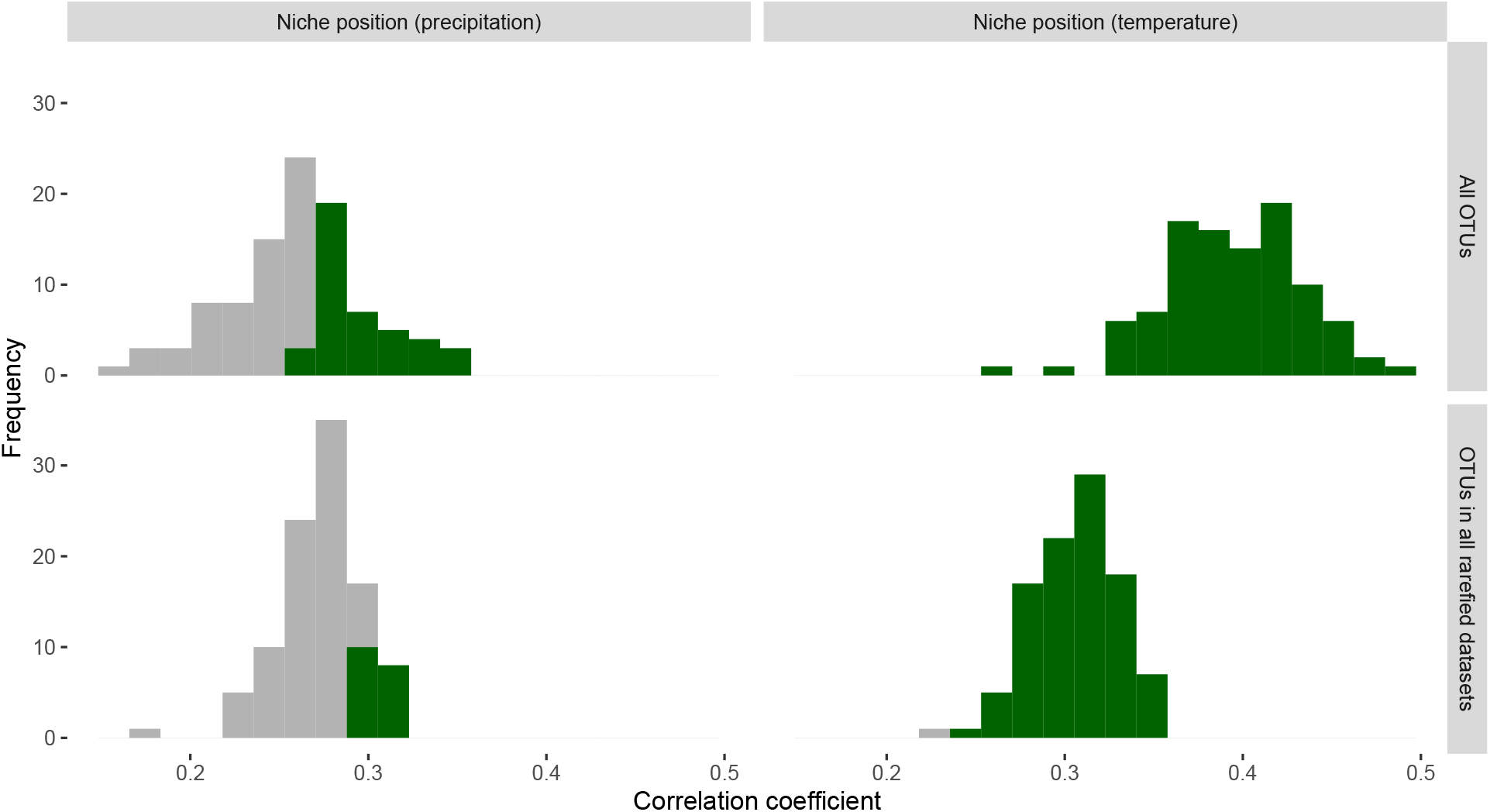
Correlation of climatic niches of arthropod OTUs between transects. Histograms represent Spearman’s correlation coefficients of climatic niche position across 100 rarefied datasets for all OTUs (top panels) and OTUs that occur in all rarefied datasets (bottom panels). Green bars represent the proportion of tests that were significant (*α*= 0.05) based on permutation of values. Analysis was performed on OTUs that occurred in at least 5 sites on each transect.

## 4 Discussion

### 4.1 Niche conservatism drives community assembly along Hawaiian mountains

Deterministic processes such as habitat filtering play a major role in community assembly (Cavender-Bares *et al.*, 2009). Using a large-scale metabarcoding dataset, we show that climatic factors strongly shape turnover in arthropod communities, both within and between two mountain transects along Mauna Kea (Laupāhoehoe) and Mauna Loa (Stainback) on Hawai‘i Island. Specifically, we find that OTUs showed highly similar temperature niche position between mountains, suggesting that niche conservatism plays an important role in shaping the taxonomic composition of communities during colonization. Strong temperature gradients may thus represent a barrier that many lowland taxa are unlikely to overcome through natural selection, at least not as quickly as community assembly may be accomplished through dispersal from other, potentially distant, highland sources (Merckx *et al.*, 2015). These findings generally confirm our first hypothesis, and are largely consistent with previous work which has shown that strong environmental conservatism dominates the adaptive radiation of insects on oceanic islands, in which adaptive radiation has occurred largely within a given environmental envelope (Hiller *et al.*, 2019; Dorey *et al.*, 2020). We found that the no niche positions of OTU for rainfall were not strongly correlated between transects. This is perhaps due to differences in the way precipitation changes with elevation between the two transects (Supplementary Fig. S1), with Stainback exhibiting a marked increase in precipitation above 1300 m. This may also suggest that most arthropods are probably tolerant of wide ranges in precipitation.

### 4.2 Elevation isolates arthropod communities between Hawaiian mountains

While we find that niche conservatism influences dispersal between different elevation bands on the two mountains, we also find that communities are more similar within a mountain than between mountains, suggesting that arthropod communities on each mountain eventually develop their own regional pool of species. Specifically, and in support of our second hypothesis, we find that communities between Laupāhoehoe and Stainback appear increasingly different at higher elevation, further supporting the role of climate and topography in influencing diversification and diversity patterns. Mountain tops have long been hypothesized to act as species pumps (e.g., Mayr 1947; Janzen 1967; Ghalambor *et al.* 2006; Steinbauer *et al.* 2016; Rahbek *et al.* 2019b), with climatic oscillations and topographic complexity causing repeated admixture and isolation events that may promote speciation (Gillespie & Roderick, 2014; Gillespie *et al.*, 2020; Rahbek *et al.*, 2019a; Salces-Castellano *et al.*, 2020). This topography-driven speciation model is dependent on the conservation of niches within lineages (Wiens, 2004; Wiens & Graham, 2005). It is possible that dispersal filtering may play a role in shaping the pattern of increasing community dissimilarity with elevation. For example, high elevations may be compositionally different between transects simply because highlands host taxa with poor dispersal abilities. However, we argue that this to be unlikely or play a minor role in shaping our findings. The observed patterns were largely consistent across different orders of arthropods, which comprise ecologically very diverse groups with various dispersal capabilities and ecologies (predators, herbivores and detritivores). It should be noted, however, that our data does not exclude the possibility of in-situ niche differentiation after colonization, (e.g., through adaptation at range boundaries). If colonization between transects was purely based on niche conservatism, fewer taxa should have colonized the higher elevations of Mauna Loa than the lower ones. This should be reflected in a lower species diversity with increasing elevation. Only rapid speciation at the young mountain top or colonization and possibly adaptation of additional low elevation taxa could explain the equally high diversity of low and high elevations.

Furthermore, the observed pattern of increasing differentiation with elevation is also mirrored in the haplotypic variation within OTUs. Specifically, OTUs tend to be share fewer zOTUs at higher elevations, suggesting that shared taxa between the transects showed less genetic connectivity at higher elevations. One explanation for this is that the isolation of high elevation environments has promoted genetic differentiation between populations on both mountains, a phenomenon that is less pronounced at lower elevations. Alternatively, genetic differentation would have been promoted by bottlenecks during colonization of Mauna Loa from Mauna Kea populations, but we argue that such as effect would have also influenced the haplotype variation of lowland sites. Thus, the low frequency of dispersal and admixture between high elevation sites relative to low elevation areas is a more parsimonious explanation of such patterns.

The elevational range we sampled does not cover the whole distribution of wet forest along Hawai‘i’s eastern flank (approximately 1,000 m elevation). However, despite the short elevational range sampled, we still discovered a very strong pattern of species turnover (Fig. 3a). Future work should expand the gradients towards higher elevations, in particular above the trade wind inversion (ca. 2000m), where habitats shift abruptly from wet/mesic to dry/arid (Vorsino *et al.*, 2014). Another interesting addition would be the sampling of the dry Leeward sites to better disentangle the relevance of temperature versus precipitation in dictating patterns of niche conservation and species distribution. Moreover, further transects should be sampled on older Hawaiian volcanoes, to test the generality of the observed patterns, and whether the pattern of greater levels of endemicity at higher elevation is enhanced through geological time (Roderick *et al.*, 2012). For direct quantification of the effect of mountains on gene flow and in promoting speciation, future work should focus on population genetic differentiation of species within the transects (Salces-Castellano *et al.*, 2020).

### 4.3 Semi-quantitative community level metabarcoding

Our study demonstrates the utility of metabarcoding approaches to characterize ecological communities and quantify ecological variation. While amplification bias can skew the relative abundances of different taxa within a community, large scale relative abundance patterns for single taxa can be scored with this method (Amend *et al.*, 2010; Krehenwinkel *et al.*, 2017). To make species abundances from community samples comparable, specimen numbers have to be incorporated in the rarefaction process (Vandeputte *et al.*, 2017). Additional size sorting of specimen prior to sequencing enabled us to mitigate the inherent bias introduced by large taxa. The majority of Hawaiian arthropod taxa are fairly small, making four size categories sufficient. In different ecosystems, with more size variation of specimens, additional categories may have to be introduced.

As an alternative to size sorting, subsampling of tissues from each specimen could also help reduce bias. However, this will come with a considerably increased effort. In future metabarcoding studies, size informed relative abundance measures could be used for large scale analysis of species distribution patterns and niche occupancy, offering an exciting new perspective of metabarcoding from simple scoring of presence and absence of taxa.

## 5 Conclusions

Mountains are crucibles for the formation of new species and are host to some of the most unique ecosystems on earth. In this study, we characterize changes in arthropod communities along elevation gradients on the island of Hawai‘i using a novel metabarcoding approach. We show that climate is an important dimension by which arthropod communities assemble and differentiate over space and time and that mountains serve as efficient barriers to dispersal and gene flow. We hope our study advances the frontiers of metabarcoding and their utility in community-level studies.

## Acknowledgments

HK was supported by a postdoctoral fellowship of the German Research Foundation (DFG). JYL was supported by the NTU Presidential Postdoctoral Fellowship. SN was supported by a postdoctoral fellowship of the Japan Society for the Promotion of Science (JSPS). JP was supported by the MICINN through the Ramón y Cajal Program (RYC-2016-20506), and the European Union through a Marie Sklodowska-Curie COFUND, Researchers’ Night and Individual Fellowships Global (MSCA grant agreement No 747238, ‘UNISLAND’). We also thank Guilemette de Kerdrel, Savannah Miller, Edward Greg Huang and Susan Kennedy for their help sampling the arthropod specimens. For access to field sites we would like to acknowledge support and assistance from the following: Steve Bergfeld (Department of Forestry and Wildlife, DOFAW), Tabetha Block (Hawai‘i Experimental Tropical Forest, HETF), Charmian Dang (Natural Areas Reserve System, NARS), Melissa Dean (HETF), Betsy Gagne (NAR), Cynthia King (Department of Land and Natural Resources, DLNR), and Joey Mello (DOFAW). The research was supported by a NSF Dimensions in Biodiversity grant (DEB 1241253) to RGG.

## Data availability statement

Raw reads, OTU tables including specimen counts are available as a Dryad digital repository (doi:10.5061/dryad.wdbrv15p5).

## Author contributions

HK, SN, and LC collected the specimens and performed size sorting and counting. HK and SN performed molecular processing of the samples. JYL, HK and JP analyzed the data. JYL, HK, JP and RG wrote the manuscript with input and comments for all other co-authors.

## References

Altschul, S.F., Gish, W., Miller, W., Myers, E.W. & Lipman, D.J.(1990) Basic local alignment search tool. Journal of Molecular Biology, 215, 403–410.

Amend, A.S., Seifert, K.A. & Bruns, T.D.(2010) Quantifying microbial communities with 454 pyrosequencing: does read abundance count? Molecular Ecology, 19, 5555–5565.

Burnham, K.P. & Anderson, D.R.(2002) Model selection and multimodel inference: A practical information-theoretic approach. Springer New York.

Camacho-Sanchez, M., Hawkins, M.T., Yu, F.T.Y., Maldonado, J.E. & Leonard, J.A. (2019) Endemism and diversity of small mammals along two neighboring bornean mountains. PeerJ, 7, e7858.

Carson, H.L., Lockwood, J.P. & Craddock, E.M.(1990) Extinction and recolonization of local populations on a growing shield volcano. Proceedings of the National Academy of Sciences, 87, 7055–7057.

Cavender-Bares, J., Kozak, K.H., Fine, P.V. & Kembel, S.W. (2009) The merging of community ecology and phylogenetic biology. Ecology Letters, 12, 693–715.

de Kerdrel, G.A., Andersen, J.C., Kennedy, S.R., Gillespie, R. & Krehenwinkel, H. (2020) Rapid and cost-effective generation of single specimen multilocus barcoding data from whole arthropod communities by multiple levels of multiplexing. Scientific Reports, 10, 1–12.

De Meester, L., Vanoverbeke, J., Kilsdonk, L.J. & Urban, M.C. (2016) Evolving perspectives on monopolization and priority effects. Trends in Ecology & Evolution, 31, 136–146.

Dorey, J.B., Groom, S.V., Freedman, E.H., Matthews, C.S., Davies, O.K., Deans, E.J., Rebola, C., Stevens, M.I., Lee, M.S. & Schwarz, M.P.(2020) Radiation of tropical island bees and the role of phylogenetic niche conservatism as an important driver of biodiversity. Proceedings of the Royal Society B, 287, 20200045.

Edgar, R.C. (2010) Search and clustering orders of magnitude faster than BLAST. Bioinformatics, 26, 2460–2461.

Edgar, R.C. (2016) UNOISE2: improved error-correction for Illumina 16S and ITS amplicon sequencing. BioRxiv, 081257.

Ghalambor, C.K., Huey, R.B., Martin, P.R., Tewksbury, J.J. & Wang, G.(2006) Are mountain passes higher in the tropics? Janzen’s hypothesis revisited. Integrative and Comparative Biology, 46, 5–17.

Giambelluca, T., Shuai, X., Barnes, M.L., Alliss, R.J., Longman, R.J., Miura, T., Chen, Q., Frazier, A.G., Mudd, R.G., Cuo, L. & Businger, A.D. (2014) Evapotranspiration of Hawai‘i. Final report submitted to the US Army Corps of Engineers. Honolulu District, and the Commission on Water Resource Management, State of Hawai ‘i.

Giambelluca, T.W., Chen, Q., Frazier, A.G., Price, J.P., Chen, Y.L., Chu, P.S., Eischeid, J.K. & Delparte, D.M.(2013) Online rainfall atlas of Hawai‘i. Bulletin of the American Meteorological Society, 94, 313–316.

Gibson, J., Shokralla, S., Porter, T.M., King, I., van Konynenburg, S., Janzen, D.H., Hallwachs, W. & Hajibabaei, M. (2014) Simultaneous assessment of the macrobiome and microbiome in a bulk sample of tropical arthropods through dna metasystematics. Proceedings of the National Academy of Sciences, 111, 8007–8012.

Gillespie, R.G., Lim, J.Y. & Rominger, A.J.(2020) The theory of evolutionary biogeography. S.M. Scheiner & D.P. Mindell, eds., The Theory of Evolution, The University of Chicago Press Chicago, IL, pp. 319–337.

Gillespie, R.G. & Roderick, G.K.(2014) Geology and climate drive diversification. Nature, 509, 297–298.

Giribet, G. & Edgecombe, G.D. (2012) Reevaluating the arthropod tree of life. Annual Review of Entomology, 57, 167–186.

Gordon, A. & Hannon, G.J. (2010) Fastx-toolkit. FASTQ/A short-reads preprocessing tools (unpublished) http://hannonlab.cshl.edu/fastx_toolkit, 5.

Halsch, C.A., Shapiro, A.M., Fordyce, J.A., Nice, C.C., Thorne, J.H., Waetjen, D.P. & Forister, M.L.(2021) Insects and recent climate change. Proceedings of the National Academy of Sciences, 118.

Heaney, L.R. (2001) Small mammal diversity along elevational gradients in the philippines: an assessment of patterns and hypotheses. Global Ecology and Biogeography, 10, 15–39.

Hiller, A.E., Koo, M.S., Goodman, K.R., Shaw, K.L., O’Grady, P.M. & Gillespie, R.G. (2019) Niche conservatism predominates in adaptive radiation: comparing the diversification of hawaiian arthropods using ecological niche modelling. Biological Journal of the Linnean Society, 127, 479–492.

Hubbell, S.P. (2001) The unified neutral theory of biodiversity and biogeography. Princeton University Press, Princeton, New Jersey, U.S.A.

Irl, S.D., Harter, D.E., Steinbauer, M.J., Gallego Puyol, D., Fernández-Palacios, J.M., Jentsch, A. & Beierkuhnlein,C. (2015) Climate vs. topography–spatial patterns of plant species diversity and endemism on a high-elevation island. Journal of Ecology, 103, 1621–1633.

Janzen, D.H. (1967) Why mountain passes are higher in the tropics. The American Naturalist, 101, 233–249.

Kessler, M. & Kluge, J. (2008) Diversity and endemism in tropical montane forests-from patterns to processes. S.R. Gradstein, J. Homeier & G. D, eds., The Tropical Mountain Forest - Patterns and Processes in a Biodiversity Hotspot, Universitötsverlag Göttingen, Göttingen, Germany, Göttingen, Germany, pp. 35–50.

Kivimäki, I., Shimbo, M. & Saerens, M. (2014) Developments in the theory of randomized shortest paths with a comparison of graph node distances. Physica A: Statistical Mechanics and its Applications, 393, 600–616.

Kozak, K.H. & Wiens, J.J.(2010) Accelerated rates of climatic-niche evolution underlie rapid species diversification. Ecology Letters, 13, 1378–1389.

Krehenwinkel, H., Wolf, M., Lim, J.Y., Rominger, A.J., Simison, W.B. & Gillespie, R.G. (2017) Estimating and mitigating amplification bias in qualitative and quantitative arthropod metabarcoding. Scientific Reports, 7, 1–12.

Lange, V., Böhme, I., Hofmann, J., Lang, K., Sauter, J., Schöne, B., Paul, P., Albrecht, V., Andreas, J.M., Baier, D.M., Nething, J., Ehninger, U., Schwarzelt, C., Pingel, J., Ehninger, G. & Schmidt, A.H. (2014) Cost-efficient high-throughput hla typing by miseq amplicon sequencing. BMC Genomics, 15, 1–11.

Leray, M., Yang, J.Y., Meyer, C.P., Mills, S.C., Agudelo, N., Ranwez, V., Boehm, J.T. & Machida, R.J. (2013) A new versatile primer set targeting a short fragment of the mitochondrial COI region for metabarcoding metazoan diversity: application for characterizing coral reef fish gut contents. Frontiers in Zoology, 10, 1–14.

Macedo, M.V., Monteiro, R.F., Flinte, V., Almeida-Neto, M., Khattar, G., da Silveira, L.F.L., de O Araujo, C., de O Araujo, R., Colares, C., da Gomes, C.V.S., Mendes, C.B., Santos, E.F. & Mayhew, P.J. (2018) Insect elevational specialization in a tropical biodiversity hotspot. Insect Conservation and Diversity, 11, 240–254.

Mayr, E. (1947) Ecological factors in speciation. Evolution, 1, 263–288.

McRae, B.H., Dickson, B.G., Keitt, T.H. & Shah, V.B.(2008) Using circuit theory to model connectivity in ecology, evolution, and conservation. Ecology, 89, 2712–2724.

Merckx, V.S.F.T., Hendriks, K.P., Beentjes, K.K., Mennes, C.B., Becking, L.E., Peijnenburg, K.T.C.A., Afendy, A., Arumugam, N., de Boer, H., Biun, A., Buang, M.M., Chen, P.P., Chung, A.Y.C., Dow, R., Feijen, F.A.A., Feijen, H., Soest, C.F.v., Geml, J., Geurts, R., Gravendeel, B., Hovenkamp, P., Imbun, P., Ipor, I., Janssens, S.B., Jocqué, M., Kappes, H., Khoo, E., Koomen, P., Lens, F., Majapun, R.J., Morgado, L.N., Neupane, S., Nieser, N., Pereira, J.T., Rahman, H., Sabran, S., Sawang, A., Schwallier, R.M., Shim, P.S., Smit, H., Sol, N., Spait, M., Stech, M., Stokvis, F., Sugau, J.B., Suleiman, M., Sumail, S., Thomas, D.C., van Tol, J., Tuh, F.Y.Y., Yahya, B.E., Nais, J., Repin, R., Lakim, M. & Schilthuizen, M. (2015) Evolution of endemism on a young tropical mountain. Nature, 524, 347–350.

Oksanen, J., Blanchet, F.G., Friendly, M., Kindt, R., Legendre, P., McGlinn, D., Minchin, P.R., O’Hara, R.B., Simpson, G.L., Solymos, P., Stevens, M.H.H., Szoecs, E. & Wagner, H. (2020) vegan: Community Ecology Package. R package version 2.5-7.

Peterman, W.E. (2018) ResistanceGA: An R package for the optimization of resistance surfaces using genetic algorithms. Methods in Ecology and Evolution, 9, 1638–1647.

Peterman, W.E., Connette, G.M., Semlitsch, R.D. & Eggert, L.S.(2014) Ecological resistance surfaces predict fine-scale genetic differentiation in a terrestrial woodland salamander. Molecular Ecology, 23, 2402–2413.

Price, J.P. (2004) Floristic biogeography of the hawaiian islands: influences of area, environment and paleogeography. Journal of Biogeography, 31, 487–500.

Rahbek, C., Borregaard, M.K., Antonelli, A., Colwell, R.K., Holt, B.G., Nogues-Bravo, D., Rasmussen, C.M., Richardson, K., Rosing, M.T., Whittaker, R.J. & Fjeldså, J. (2019a) Building mountain biodiversity: geological and evolutionary processes. Science, 365, 1114–1119.

Rahbek, C., Borregaard, M.K., Colwell, R.K., Dalsgaard, B., Holt, B.G., Morueta-Holme, N., Nogues-Bravo, D., Whittaker, R.J. & Fjeldså, J. (2019b) Humboldt’s enigma: What causes global patterns of mountain biodiversity? Science, 365, 1108–1113.

Ricklefs, R.E. (2004) A comprehensive framework for global patterns in biodiversity. Ecology Letters, 7, 1–15.

Roderick, G.K., Croucher, P.J., Vandergast, A.G. & Gillespie, R.G.(2012) Species differentiation on a dynamic landscape: shifts in metapopulation genetic structure using the chronology of the hawaiian archipelago. Evolutionary Biology, 39, 192–206.

Rominger, A.J., Goodman, K.R., Lim, J.Y., Armstrong, E.E., Becking, L.E., Bennett, G.M., Brewer, M.S., Cotoras, D.D., Ewing, C.P., Harte, J., Martinez, N.D., O’Grady, P.M., Percy, D.M., Price, D.K., Roderick, G.R., Shaw, K.L., Valdovinos, F.S., Gruner, D.S. & Gillespie, R.G. (2016) Community assembly on isolated islands: macroecology meets evolution. Global Ecology and Biogeography, 25, 769–780.

Row, J.R., Knick, S.T., Oyler-McCance, S.J., Lougheed, S.C. & Fedy, B.C. (2017) Developing approaches for linear mixed modeling in landscape genetics through landscape-directed dispersal simulations. Ecology and Evolution, 7, 3751–3761.

Salces-Castellano, A., Patiño, J., Alvarez, N., Andújar, C., Arribas, P., Braojos-Ruiz, J.J., del Arco-Aguilar, M., García-Olivares, V., Karger, D.N., López, H. et al. (2020) Climate drives community-wide divergence within species over a limited spatial scale: evidence from an oceanic island. Ecology Letters, 23, 305–315.

Shaw, K.L. & Gillespie, R.G.(2016) Comparative phylogeography of oceanic archipelagos: Hotspots for inferences of evolutionary process. Proceedings of the National Academy of Sciences, 113, 7986–7993.

Sherrod, D.R., Sinton, J.M., Watkins, S.E. & Brunt, K.M.(2007) Geologic Map of the State of Hawai‘i. US Geological Survey Open-File Report 2007-1089.

Soberón, J. (2007) Grinnellian and Eltonian niches and geographic distributions of species. Ecology Letters, 10, 1115–1123.

Steinbauer, M.J., Field, R., Grytnes, J.A., Trigas, P., Ah-Peng, C., Attorre, F., Birks, H.J.B., Borges, P.A., Cardoso, P., Chou, C.H., Sanctis, M.D., de Sequeira, M.M., Duarte, M.C., Elias, R.B., Fernández-Palacios, J.M., Gabriel, R., Gereau, R.E., Gillespie, R.G., Greimler, J., Harter, D.E.V., Huang, T.J., Irl, S.D.H., Jeanmonod, D., Jentsch, A., Jump, A.S., Kueffer, C., Nogué, S., Otto, R., Price, J., Romeiras, M.M., Strasberg, D., Stuessy, T., Svenning, J.C., Vetass, O.R. & Beierkuhnlein, C. (2016) Topography-driven isolation, speciation and a global increase of endemism with elevation. Global Ecology and Biogeography, 25, 1097–1107.

Tamura, K., Stecher, G., Peterson, D., Filipski, A. & Kumar, S. (2013) MEGA6: molecular evolutionary genetics analysis version 6.0. Molecular Biology and Evolution, 30, 2725–2729.

van Etten, J. (2017) R package gdistance: distances and routes on geographical grids. Journal of Statistical Software, 76, 1–21.

Vanbergen, A.J. & the Insect Pollinators Initiative (2013) Threats to an ecosystem service: pressures on pollinators. Frontiers in Ecology and the Environment, 11, 251–259.

Vandeputte, D., Kathagen, G., D’hoe, K., Vieira-Silva, S., Valles-Colomer, M., Sabino, J., Wang, J., Tito, R.Y., De Commer, L., Darzi, Y. et al. (2017) Quantitative microbiome profiling links gut community variation to microbial load. Nature, 551, 507–511.

Vasconcelos, T.N., Alcantara, S., Andrino, C.O., Forest, F., Reginato, M., Simon, M.F. & Pirani, J.R. (2020) Fast diversification through a mosaic of evolutionary histories characterizes the endemic flora of ancient neotropical mountains. Proceedings of the Royal Society B, 287, 20192933.

Vellend, M. (2016) The theory of ecological communities. Princeton University Press, Princeton, New Jersey, U.S.A.

Vorsino, A.E., Fortini, L.B., Amidon, F.A., Miller, S.E., Jacobi, J.D., Price, J.P., Koob, G.A. et al. (2014) Modeling hawaiian ecosystem degradation due to invasive plants under current and future climates. PloS one, 9, e95427.

Webb, C.O., Ackerly, D.D., McPeek, M.A. & Donoghue, M.J.(2002) Phylogenies and community ecology. Annual Review of Ecology and Systematics, 33, 475–505.

Wiens, J.J. (2004) Speciation and ecology revisited: phylogenetic niche conservatism and the origin of species. Evolution, 58, 193–197.

Wiens, J.J. & Graham, C.H.(2005) Niche conservatism: integrating evolution, ecology, and conservation biology. Annual Review of Ecology, Evolution and Systematics, 36, 519–539.

Yoder, J.B., Clancey, E., Des Roches, S., Eastman, J., Gentry, L., Godsoe, W., Hagey, T., Jochimsen, D., Oswald, B.P., Robertson, J., Sarver, B.A.J., Schenk, J.J., Spear, S.F. & Harmon, L.J. (2010) Ecological opportunity and the origin of adaptive radiations. Journal of Evolutionary Biology, 23, 1581–1596.

Zhang, J., Kobert, K., Flouri, T. & Stamatakis, A. (2014) PEAR: a fast and accurate Illumina Paired-End reAd mergeR. Bioinformatics, 30, 614–620.

