## Supplementary Material for "Climatic niche conservatism shapes the ecological assembly of Hawaiian arthropod communities"

**Table of Contents**

- Supplementary Table 1
- Supplementary Figures 1 – 6

### Supplementary Tables

**Supplementary Table 1: Akaike weights (w), ΔAICc, and marginal ($R^2{}_m$) and conditional ($R^2{}_c$) R-square values for resistance models.** Models were fit for each taxonomic order on each transect separately (L = Laupāhoehoe, S = Stainback). Analyses were implemented for five randomly chosen rarefied datasets.

| Rarefied_ID | Transect | Order | Model | $R^2{}_m$ | $R^2{}_c$ | ΔAICc | w |
|---|---|---|---|---|---|---|---|
| 14 | L | Araneae | Geographic distance | 0.113 | 0.285 | 0 | 0.72 |
| 14 | L | Araneae | Precip | 0.13 | 0.27 | 3.104 | 0.153 |
| 14 | L | Araneae | Temp | 0.125 | 0.273 | 3.602 | 0.119 |
| 14 | L | Araneae | Temp + Precip | 0.113 | 0.285 | 8.929 | 0.008 |
| 14 | L | Araneae | Null | 0 | 0.143 | 28.74 | <0.01 |
| 51 | L | Araneae | Geographic distance | 0.115 | 0.281 | 0 | 0.729 |
| 51 | L | Araneae | Precip | 0.129 | 0.268 | 3.235 | 0.145 |
| 51 | L | Araneae | Temp | 0.126 | 0.27 | 3.637 | 0.118 |
| 51 | L | Araneae | Temp + Precip | 0.115 | 0.281 | 8.929 | 0.008 |
| 51 | L | Araneae | Null | 0 | 0.138 | 29.188 | <0.01 |
| 80 | L | Araneae | Geographic distance | 0.117 | 0.273 | 0 | 0.762 |
| 80 | L | Araneae | Precip | 0.124 | 0.26 | 3.638 | 0.124 |
| 80 | L | Araneae | Temp | 0.126 | 0.263 | 3.947 | 0.106 |
| 80 | L | Araneae | Temp + Precip | 0.117 | 0.273 | 8.929 | 0.009 |
| 80 | L | Araneae | Null | 0 | 0.129 | 29.311 | <0.01 |
| 90 | L | Araneae | Geographic distance | 0.108 | 0.284 | 0 | 0.716 |
| 90 | L | Araneae | Precip | 0.118 | 0.268 | 3.086 | 0.153 |
| 90 | L | Araneae | Temp | 0.12 | 0.273 | 3.522 | 0.123 |
| 90 | L | Araneae | Temp + Precip | 0.108 | 0.284 | 8.929 | 0.008 |
| 90 | L | Araneae | Null | 0 | 0.141 | 27.164 | <0.01 |
| 92 | L | Araneae | Geographic distance | 0.111 | 0.305 | 0 | 0.785 |
| 92 | L | Araneae | Precip | 0.119 | 0.288 | 3.914 | 0.111 |
| 92 | L | Araneae | Temp | 0.119 | 0.293 | 4.235 | 0.095 |
| 92 | L | Araneae | Temp + Precip | 0.111 | 0.305 | 8.929 | 0.009 |
| 92 | L | Araneae | Null | 0 | 0.169 | 29.005 | <0.01 |

| | | | | | | | |
|---|---|---|---|---|---|---|---|
| 14 | S | Araneae | Temp | 0.696 | 0.867 | 0 | 0.798 |
| 14 | S | Araneae | Temp + Precip | 0.671 | 0.828 | 2.754 | 0.201 |
| 14 | S | Araneae | Precip | 0.715 | 0.919 | 21.563 | <0.01 |
| 14 | S | Araneae | Geographic distance | 0.324 | 0.61 | 99.614 | <0.01 |
| 14 | S | Araneae | Null | 0 | 0.314 | 455.857 | <0.01 |
| 51 | S | Araneae | Temp | 0.68 | 0.833 | 0 | 0.688 |
| 51 | S | Araneae | Temp + Precip | 0.683 | 0.845 | 1.579 | 0.312 |
| 51 | S | Araneae | Precip | 0.603 | 0.838 | 23.165 | <0.01 |
| 51 | S | Araneae | Geographic distance | 0.34 | 0.612 | 97.961 | <0.01 |
| 51 | S | Araneae | Null | 0 | 0.299 | 468.961 | <0.01 |
| 80 | S | Araneae | Precip | 0.629 | 0.775 | 0 | 0.952 |
| 80 | S | Araneae | Temp | 0.708 | 0.904 | 5.988 | 0.048 |
| 80 | S | Araneae | Temp + Precip | 0.559 | 0.77 | 19.449 | <0.01 |
| 80 | S | Araneae | Geographic distance | 0.326 | 0.592 | 90.329 | <0.01 |
| 80 | S | Araneae | Null | 0 | 0.294 | 436.256 | <0.01 |
| 90 | S | Araneae | Precip | 0.676 | 0.836 | 0 | 0.912 |
| 90 | S | Araneae | Temp + Precip | 0.674 | 0.834 | 4.903 | 0.079 |
| 90 | S | Araneae | Temp | 0.651 | 0.833 | 9.119 | 0.01 |
| 90 | S | Araneae | Geographic distance | 0.335 | 0.594 | 93.139 | <0.01 |
| 90 | S | Araneae | Null | 0 | 0.285 | 448.439 | <0.01 |
| 92 | S | Araneae | Precip | 0.697 | 0.862 | 0 | 0.701 |
| 92 | S | Araneae | Temp + Precip | 0.672 | 0.836 | 2.836 | 0.17 |
| 92 | S | Araneae | Temp | 0.67 | 0.826 | 3.371 | 0.13 |
| 92 | S | Araneae | Geographic distance | 0.342 | 0.614 | 96.356 | <0.01 |
| 92 | S | Araneae | Null | 0 | 0.299 | 471.151 | <0.01 |
| 14 | L | Coleoptera | Temp | 0.272 | 0.455 | 0 | 0.541 |
| 14 | L | Coleoptera | Precip | 0.252 | 0.446 | 0.512 | 0.419 |
| 14 | L | Coleoptera | Geographic distance | 0.133 | 0.425 | 5.217 | 0.04 |
| 14 | L | Coleoptera | Temp + Precip | 0.133 | 0.425 | 14.146 | <0.01 |
| 14 | L | Coleoptera | Null | 0 | 0.286 | 47.116 | <0.01 |
| 51 | L | Coleoptera | Temp | 0.285 | 0.463 | 0 | 0.492 |
| 51 | L | Coleoptera | Precip | 0.259 | 0.448 | 0.147 | 0.457 |

| | | | | | | | |
|---|---|---|---|---|---|---|---|
| 51 | L | Coleoptera | Geographic distance | 0.163 | 0.426 | 4.533 | 0.051 |
| 51 | L | Coleoptera | Temp + Precip | 0.163 | 0.426 | 13.462 | 0.001 |
| 51 | L | Coleoptera | Null | 0 | 0.273 | 55.929 | <0.01 |
| 80 | L | Coleoptera | Temp | 0.231 | 0.444 | 0 | 0.428 |
| 80 | L | Coleoptera | Precip | 0.224 | 0.443 | 0.268 | 0.374 |
| 80 | L | Coleoptera | Geographic distance | 0.131 | 0.426 | 2.566 | 0.196 |
| 80 | L | Coleoptera | Temp + Precip | 0.131 | 0.426 | 10.495 | 0.002 |
| 80 | L | Coleoptera | Null | 0 | 0.285 | 42.881 | <0.01 |
| 90 | L | Coleoptera | Temp | 0.262 | 0.466 | 0 | 0.44 |
| 90 | L | Coleoptera | Precip | 0.239 | 0.456 | 0.114 | 0.415 |
| 90 | L | Coleoptera | Geographic distance | 0.162 | 0.449 | 2.245 | 0.143 |
| 90 | L | Coleoptera | Temp + Precip | 0.162 | 0.449 | 11.174 | 0.002 |
| 90 | L | Coleoptera | Null | 0 | 0.283 | 55.029 | <0.01 |
| 92 | L | Coleoptera | Temp | 0.274 | 0.486 | 0 | 0.517 |
| 92 | L | Coleoptera | Precip | 0.243 | 0.47 | 0.434 | 0.416 |
| 92 | L | Coleoptera | Geographic distance | 0.149 | 0.46 | 4.096 | 0.067 |
| 92 | L | Coleoptera | Temp + Precip | 0.149 | 0.46 | 13.026 | 0.001 |
| 92 | L | Coleoptera | Null | 0 | 0.305 | 53.61 | <0.01 |
| 14 | S | Coleoptera | Temp + Precip | 0.135 | 0.185 | 0 | 0.701 |
| 14 | S | Coleoptera | Precip | 0.104 | 0.171 | 3.051 | 0.153 |
| 14 | S | Coleoptera | Geographic distance | 0.074 | 0.159 | 4.468 | 0.075 |
| 14 | S | Coleoptera | Temp | 0.1 | 0.173 | 4.586 | 0.071 |
| 14 | S | Coleoptera | Null | 0 | 0.095 | 52.869 | <0.01 |
| 51 | S | Coleoptera | Geographic distance | 0.083 | 0.15 | 0 | 0.35 |
| 51 | S | Coleoptera | Precip | 0.102 | 0.158 | 0.239 | 0.31 |
| 51 | S | Coleoptera | Temp + Precip | 0.11 | 0.16 | 0.86 | 0.228 |
| 51 | S | Coleoptera | Temp | 0.094 | 0.156 | 2.277 | 0.112 |
| 51 | S | Coleoptera | Null | 0 | 0.075 | 53.487 | <0.01 |
| 80 | S | Coleoptera | Geographic distance | 0.077 | 0.16 | 0 | 0.455 |
| 80 | S | Coleoptera | Temp + Precip | 0.121 | 0.183 | 1.221 | 0.247 |
| 80 | S | Coleoptera | Precip | 0.092 | 0.167 | 1.776 | 0.187 |
| 80 | S | Coleoptera | Temp | 0.087 | 0.166 | 2.81 | 0.112 |

| | | | | | | | |
|---|---|---|---|---|---|---|---|
| 80 | S | Coleoptera | Null | 0 | 0.092 | 50.327 | <0.01 |
| 90 | S | Coleoptera | Geographic distance | 0.068 | 0.143 | 0 | 0.487 |
| 90 | S | Coleoptera | Temp + Precip | 0.113 | 0.17 | 1.583 | 0.221 |
| 90 | S | Coleoptera | Precip | 0.082 | 0.149 | 1.913 | 0.187 |
| 90 | S | Coleoptera | Temp | 0.076 | 0.147 | 3.077 | 0.105 |
| 90 | S | Coleoptera | Null | 0 | 0.083 | 43.421 | <0.01 |
| 92 | S | Coleoptera | Temp + Precip | 0.141 | 0.191 | 0 | 0.69 |
| 92 | S | Coleoptera | Precip | 0.113 | 0.179 | 3.052 | 0.15 |
| 92 | S | Coleoptera | Temp | 0.099 | 0.169 | 4.307 | 0.08 |
| 92 | S | Coleoptera | Geographic distance | 0.078 | 0.159 | 4.316 | 0.08 |
| 92 | S | Coleoptera | Null | 0 | 0.092 | 54.956 | <0.01 |
| 14 | L | Hemiptera | Precip | 0.312 | 0.422 | 0 | 0.437 |
| 14 | L | Hemiptera | Temp | 0.306 | 0.416 | 0.422 | 0.354 |
| 14 | L | Hemiptera | Geographic distance | 0.213 | 0.358 | 1.493 | 0.207 |
| 14 | L | Hemiptera | Temp + Precip | 0.213 | 0.358 | 10.423 | 0.002 |
| 14 | L | Hemiptera | Null | 0 | 0.199 | 62.215 | <0.01 |
| 51 | L | Hemiptera | Precip | 0.294 | 0.408 | 0 | 0.4 |
| 51 | L | Hemiptera | Temp | 0.288 | 0.402 | 0.397 | 0.328 |
| 51 | L | Hemiptera | Geographic distance | 0.202 | 0.35 | 0.788 | 0.27 |
| 51 | L | Hemiptera | Temp + Precip | 0.202 | 0.35 | 9.717 | 0.003 |
| 51 | L | Hemiptera | Null | 0 | 0.199 | 57.802 | <0.01 |
| 80 | L | Hemiptera | Precip | 0.294 | 0.407 | 0 | 0.398 |
| 80 | L | Hemiptera | Temp | 0.288 | 0.402 | 0.447 | 0.318 |
| 80 | L | Hemiptera | Geographic distance | 0.202 | 0.348 | 0.695 | 0.281 |
| 80 | L | Hemiptera | Temp + Precip | 0.202 | 0.348 | 9.624 | 0.003 |
| 80 | L | Hemiptera | Null | 0 | 0.198 | 57.722 | <0.01 |
| 90 | L | Hemiptera | Precip | 0.305 | 0.415 | 0 | 0.421 |
| 90 | L | Hemiptera | Temp | 0.299 | 0.409 | 0.406 | 0.343 |
| 90 | L | Hemiptera | Geographic distance | 0.208 | 0.354 | 1.183 | 0.233 |
| 90 | L | Hemiptera | Temp + Precip | 0.208 | 0.354 | 10.113 | 0.003 |
| 90 | L | Hemiptera | Null | 0 | 0.2 | 60.186 | <0.01 |
| 92 | L | Hemiptera | Precip | 0.294 | 0.403 | 0 | 0.373 |

| | | | | | | | |
|---|---|---|---|---|---|---|---|
| 92 | L | Hemiptera | Temp | 0.29 | 0.398 | 0.286 | 0.323 |
| 92 | L | Hemiptera | Geographic distance | 0.208 | 0.345 | 0.431 | 0.301 |
| 92 | L | Hemiptera | Temp + Precip | 0.208 | 0.345 | 9.361 | 0.003 |
| 92 | L | Hemiptera | Null | 0 | 0.188 | 58.853 | <0.01 |
| 14 | S | Hemiptera | Temp + Precip | 0.416 | 0.585 | 0 | 1 |
| 14 | S | Hemiptera | Temp | 0.288 | 0.502 | 21.739 | <0.01 |
| 14 | S | Hemiptera | Geographic distance | 0.189 | 0.47 | 29.306 | <0.01 |
| 14 | S | Hemiptera | Precip | 0.314 | 0.656 | 66.338 | <0.01 |
| 14 | S | Hemiptera | Null | 0 | 0.296 | 207.97 | <0.01 |
| 51 | S | Hemiptera | Temp + Precip | 0.399 | 0.562 | 0 | 0.998 |
| 51 | S | Hemiptera | Temp | 0.301 | 0.497 | 12.766 | 0.002 |
| 51 | S | Hemiptera | Precip | 0.259 | 0.48 | 21.631 | <0.01 |
| 51 | S | Hemiptera | Geographic distance | 0.182 | 0.447 | 23.392 | <0.01 |
| 51 | S | Hemiptera | Null | 0 | 0.28 | 189.6 | <0.01 |
| 80 | S | Hemiptera | Temp + Precip | 0.402 | 0.561 | 0 | 0.998 |
| 80 | S | Hemiptera | Temp | 0.279 | 0.476 | 12.775 | 0.002 |
| 80 | S | Hemiptera | Geographic distance | 0.179 | 0.44 | 21.662 | <0.01 |
| 80 | S | Hemiptera | Precip | 0.231 | 0.466 | 26.327 | <0.01 |
| 80 | S | Hemiptera | Null | 0 | 0.278 | 183.627 | <0.01 |
| 90 | S | Hemiptera | Temp + Precip | 0.428 | 0.587 | 0 | 0.986 |
| 90 | S | Hemiptera | Precip | 0.375 | 0.539 | 8.455 | 0.014 |
| 90 | S | Hemiptera | Temp | 0.325 | 0.511 | 18.498 | <0.01 |
| 90 | S | Hemiptera | Geographic distance | 0.198 | 0.46 | 29.288 | <0.01 |
| 90 | S | Hemiptera | Null | 0 | 0.277 | 212.082 | <0.01 |
| 92 | S | Hemiptera | Temp + Precip | 0.432 | 0.588 | 0 | 0.567 |
| 92 | S | Hemiptera | Temp | 0.483 | 0.677 | 0.539 | 0.433 |
| 92 | S | Hemiptera | Precip | 0.255 | 0.514 | 13.712 | 0.001 |
| 92 | S | Hemiptera | Geographic distance | 0.178 | 0.441 | 15.758 | <0.01 |
| 92 | S | Hemiptera | Null | 0 | 0.278 | 176.989 | <0.01 |
| 14 | L | Lepidoptera | Geographic distance | 0.19 | 0.268 | 0 | 0.786 |
| 14 | L | Lepidoptera | Temp | 0.209 | 0.282 | 4.048 | 0.104 |
| 14 | L | Lepidoptera | Precip | 0.208 | 0.282 | 4.098 | 0.101 |

| | | | | | | | |
|---|---|---|---|---|---|---|---|
| 14 | L | Lepidoptera | Temp + Precip | 0.19 | 0.268 | 8.929 | 0.009 |
| 14 | L | Lepidoptera | Null | 0 | 0.08 | 49.465 | <0.01 |
| 51 | L | Lepidoptera | Geographic distance | 0.223 | 0.318 | 0 | 0.776 |
| 51 | L | Lepidoptera | Temp | 0.242 | 0.331 | 3.932 | 0.109 |
| 51 | L | Lepidoptera | Precip | 0.246 | 0.334 | 3.982 | 0.106 |
| 51 | L | Lepidoptera | Temp + Precip | 0.223 | 0.318 | 8.929 | 0.009 |
| 51 | L | Lepidoptera | Null | 0 | 0.107 | 60.78 | <0.01 |
| 80 | L | Lepidoptera | Geographic distance | 0.204 | 0.3 | 0 | 0.809 |
| 80 | L | Lepidoptera | Temp | 0.218 | 0.308 | 4.331 | 0.093 |
| 80 | L | Lepidoptera | Precip | 0.22 | 0.311 | 4.407 | 0.089 |
| 80 | L | Lepidoptera | Temp + Precip | 0.204 | 0.3 | 8.929 | 0.009 |
| 80 | L | Lepidoptera | Null | 0 | 0.095 | 54.213 | <0.01 |
| 90 | L | Lepidoptera | Geographic distance | 0.182 | 0.272 | 0 | 0.849 |
| 90 | L | Lepidoptera | Temp | 0.192 | 0.278 | 4.948 | 0.072 |
| 90 | L | Lepidoptera | Precip | 0.192 | 0.279 | 5.014 | 0.069 |
| 90 | L | Lepidoptera | Temp + Precip | 0.182 | 0.272 | 8.929 | 0.01 |
| 90 | L | Lepidoptera | Null | 0 | 0.1 | 47.533 | <0.01 |
| 92 | L | Lepidoptera | Geographic distance | 0.188 | 0.309 | 0 | 0.803 |
| 92 | L | Lepidoptera | Temp | 0.206 | 0.323 | 4.207 | 0.098 |
| 92 | L | Lepidoptera | Precip | 0.207 | 0.325 | 4.376 | 0.09 |
| 92 | L | Lepidoptera | Temp + Precip | 0.188 | 0.309 | 8.929 | 0.009 |
| 92 | L | Lepidoptera | Null | 0 | 0.118 | 50.48 | <0.01 |
| 14 | S | Lepidoptera | Precip | 0.058 | 0.18 | 0 | 0.362 |
| 14 | S | Lepidoptera | Temp | 0.052 | 0.174 | 0.278 | 0.315 |
| 14 | S | Lepidoptera | Geographic distance | 0.015 | 0.146 | 1.011 | 0.219 |
| 14 | S | Lepidoptera | Temp + Precip | 0.057 | 0.179 | 2.566 | 0.1 |
| 14 | S | Lepidoptera | Null | 0 | 0.128 | 9.429 | 0.003 |
| 51 | S | Lepidoptera | Precip | 0.064 | 0.184 | 0 | 0.369 |
| 51 | S | Lepidoptera | Temp | 0.063 | 0.182 | 0.095 | 0.352 |
| 51 | S | Lepidoptera | Geographic distance | 0.016 | 0.143 | 1.776 | 0.152 |
| 51 | S | Lepidoptera | Temp + Precip | 0.067 | 0.186 | 2.159 | 0.125 |
| 51 | S | Lepidoptera | Null | 0 | 0.125 | 10.795 | 0.002 |

| | | | | | | | |
|---|---|---|---|---|---|---|---|
| 80 | S | Lepidoptera | Temp | 0.079 | 0.202 | 0 | 0.424 |
| 80 | S | Lepidoptera | Precip | 0.082 | 0.207 | 0.168 | 0.39 |
| 80 | S | Lepidoptera | Temp + Precip | 0.082 | 0.207 | 2.597 | 0.116 |
| 80 | S | Lepidoptera | Geographic distance | 0.019 | 0.159 | 3.608 | 0.07 |
| 80 | S | Lepidoptera | Null | 0 | 0.137 | 14.919 | <0.01 |
| 90 | S | Lepidoptera | Temp | 0.077 | 0.196 | 0 | 0.422 |
| 90 | S | Lepidoptera | Precip | 0.084 | 0.206 | 0.217 | 0.379 |
| 90 | S | Lepidoptera | Temp + Precip | 0.084 | 0.204 | 2.272 | 0.136 |
| 90 | S | Lepidoptera | Geographic distance | 0.022 | 0.153 | 3.791 | 0.063 |
| 90 | S | Lepidoptera | Null | 0 | 0.128 | 17.056 | <0.01 |
| 92 | S | Lepidoptera | Temp | 0.067 | 0.174 | 0 | 0.44 |
| 92 | S | Lepidoptera | Precip | 0.085 | 0.197 | 1.05 | 0.26 |
| 92 | S | Lepidoptera | Geographic distance | 0.023 | 0.139 | 2.054 | 0.158 |
| 92 | S | Lepidoptera | Temp + Precip | 0.072 | 0.181 | 2.27 | 0.142 |
| 92 | S | Lepidoptera | Null | 0 | 0.113 | 15.667 | <0.01 |
| 14 | L | Orthoptera | Temp | 0.415 | 0.415 | 0 | 0.705 |
| 14 | L | Orthoptera | Precip | 0.411 | 0.411 | 1.74 | 0.295 |
| 14 | L | Orthoptera | Geographic distance | 0.308 | 0.36 | 36.096 | <0.01 |
| 14 | L | Orthoptera | Temp + Precip | 0.308 | 0.36 | 45.026 | <0.01 |
| 14 | L | Orthoptera | Null | 0 | 0.039 | 122.468 | <0.01 |
| 51 | L | Orthoptera | Temp | 0.391 | 0.391 | 0 | 0.76 |
| 51 | L | Orthoptera | Precip | 0.385 | 0.385 | 2.304 | 0.24 |
| 51 | L | Orthoptera | Geographic distance | 0.301 | 0.347 | 32.161 | <0.01 |
| 51 | L | Orthoptera | Temp + Precip | 0.301 | 0.347 | 41.09 | <0.01 |
| 51 | L | Orthoptera | Null | 0 | 0.024 | 114.961 | <0.01 |
| 80 | L | Orthoptera | Temp | 0.421 | 0.421 | 0 | 0.629 |
| 80 | L | Orthoptera | Precip | 0.419 | 0.419 | 1.054 | 0.371 |
| 80 | L | Orthoptera | Geographic distance | 0.314 | 0.374 | 36.25 | <0.01 |
| 80 | L | Orthoptera | Temp + Precip | 0.314 | 0.374 | 45.18 | <0.01 |
| 80 | L | Orthoptera | Null | 0 | 0.041 | 124.731 | <0.01 |
| 90 | L | Orthoptera | Temp | 0.433 | 0.433 | 0 | 0.819 |
| 90 | L | Orthoptera | Precip | 0.427 | 0.427 | 3.024 | 0.181 |

| | | | | | | | |
|---|---|---|---|---|---|---|---|
| 90 | L | Orthoptera | Geographic distance | 0.309 | 0.375 | 39.674 | <0.01 |
| 90 | L | Orthoptera | Temp + Precip | 0.309 | 0.375 | 48.604 | <0.01 |
| 90 | L | Orthoptera | Null | 0 | 0.054 | 127.279 | <0.01 |
| 92 | L | Orthoptera | Temp | 0.423 | 0.423 | 0 | 0.872 |
| 92 | L | Orthoptera | Precip | 0.414 | 0.414 | 3.832 | 0.128 |
| 92 | L | Orthoptera | Geographic distance | 0.319 | 0.371 | 36.985 | <0.01 |
| 92 | L | Orthoptera | Temp + Precip | 0.319 | 0.371 | 45.914 | <0.01 |
| 92 | L | Orthoptera | Null | 0 | 0.035 | 126.687 | <0.01 |
| 14 | S | Orthoptera | Temp | 0.329 | 0.525 | 0 | 0.954 |
| 14 | S | Orthoptera | Temp + Precip | 0.272 | 0.491 | 6.108 | 0.045 |
| 14 | S | Orthoptera | Precip | 0.197 | 0.504 | 13.402 | 0.001 |
| 14 | S | Orthoptera | Geographic distance | 0.076 | 0.45 | 20.222 | <0.01 |
| 14 | S | Orthoptera | Null | 0 | 0.396 | 90.367 | <0.01 |
| 51 | S | Orthoptera | Precip | 0.264 | 0.602 | 0 | 0.497 |
| 51 | S | Orthoptera | Temp + Precip | 0.259 | 0.598 | 0.037 | 0.488 |
| 51 | S | Orthoptera | Temp | 0.188 | 0.553 | 7.107 | 0.014 |
| 51 | S | Orthoptera | Geographic distance | 0.078 | 0.504 | 16.888 | <0.01 |
| 51 | S | Orthoptera | Null | 0 | 0.42 | 100.724 | <0.01 |
| 80 | S | Orthoptera | Precip | 0.259 | 0.581 | 0 | 0.604 |
| 80 | S | Orthoptera | Temp + Precip | 0.282 | 0.597 | 0.917 | 0.382 |
| 80 | S | Orthoptera | Temp | 0.223 | 0.56 | 7.424 | 0.015 |
| 80 | S | Orthoptera | Geographic distance | 0.075 | 0.479 | 18.729 | <0.01 |
| 80 | S | Orthoptera | Null | 0 | 0.398 | 96.053 | <0.01 |
| 90 | S | Orthoptera | Precip | 0.317 | 0.633 | 0 | 0.706 |
| 90 | S | Orthoptera | Temp + Precip | 0.331 | 0.643 | 1.827 | 0.283 |
| 90 | S | Orthoptera | Temp | 0.299 | 0.625 | 8.403 | 0.011 |
| 90 | S | Orthoptera | Geographic distance | 0.08 | 0.491 | 24.256 | <0.01 |
| 90 | S | Orthoptera | Null | 0 | 0.405 | 108.279 | <0.01 |
| 92 | S | Orthoptera | Precip | 0.224 | 0.593 | 0 | 0.615 |
| 92 | S | Orthoptera | Temp + Precip | 0.198 | 0.58 | 1.104 | 0.354 |
| 92 | S | Orthoptera | Temp | 0.179 | 0.568 | 6.041 | 0.03 |
| 92 | S | Orthoptera | Geographic distance | 0.063 | 0.519 | 13.12 | 0.001 |

| | | | | | | | |
|---|---|---|---|---|---|---|---|
| 92 | S | Orthoptera | Null | 0 | 0.452 | 83.499 | <0.01 |
| 14 | L | Psocoptera | Precip | 0.36 | 0.564 | 0 | 0.699 |
| 14 | L | Psocoptera | Temp | 0.262 | 0.485 | 1.774 | 0.288 |
| 14 | L | Psocoptera | Geographic distance | 0.135 | 0.428 | 7.929 | 0.013 |
| 14 | L | Psocoptera | Temp + Precip | 0.135 | 0.428 | 16.858 | <0.01 |
| 14 | L | Psocoptera | Null | 0 | 0.309 | 51.068 | <0.01 |
| 51 | L | Psocoptera | Precip | 0.315 | 0.519 | 0 | 0.721 |
| 51 | L | Psocoptera | Temp | 0.247 | 0.465 | 2.075 | 0.256 |
| 51 | L | Psocoptera | Geographic distance | 0.131 | 0.412 | 6.921 | 0.023 |
| 51 | L | Psocoptera | Temp + Precip | 0.131 | 0.412 | 15.851 | <0.01 |
| 51 | L | Psocoptera | Null | 0 | 0.298 | 47.772 | <0.01 |
| 80 | L | Psocoptera | Precip | 0.337 | 0.532 | 0 | 0.764 |
| 80 | L | Psocoptera | Temp | 0.275 | 0.483 | 2.445 | 0.225 |
| 80 | L | Psocoptera | Geographic distance | 0.148 | 0.43 | 8.384 | 0.012 |
| 80 | L | Psocoptera | Temp + Precip | 0.148 | 0.43 | 17.313 | <0.01 |
| 80 | L | Psocoptera | Null | 0 | 0.297 | 55.735 | <0.01 |
| 90 | L | Psocoptera | Precip | 0.316 | 0.537 | 0 | 0.779 |
| 90 | L | Psocoptera | Temp | 0.238 | 0.481 | 2.747 | 0.197 |
| 90 | L | Psocoptera | Geographic distance | 0.124 | 0.433 | 6.979 | 0.024 |
| 90 | L | Psocoptera | Temp + Precip | 0.124 | 0.433 | 15.909 | <0.01 |
| 90 | L | Psocoptera | Null | 0 | 0.327 | 47.032 | <0.01 |
| 92 | L | Psocoptera | Temp | 0.265 | 0.467 | 0 | 0.586 |
| 92 | L | Psocoptera | Precip | 0.248 | 0.453 | 0.875 | 0.379 |
| 92 | L | Psocoptera | Geographic distance | 0.138 | 0.415 | 5.653 | 0.035 |
| 92 | L | Psocoptera | Temp + Precip | 0.138 | 0.415 | 14.583 | <0.01 |
| 92 | L | Psocoptera | Null | 0 | 0.292 | 48.868 | <0.01 |
| 14 | S | Psocoptera | Temp | 0.542 | 0.69 | 0 | 0.775 |
| 14 | S | Psocoptera | Temp + Precip | 0.544 | 0.692 | 2.733 | 0.198 |
| 14 | S | Psocoptera | Precip | 0.533 | 0.69 | 6.713 | 0.027 |
| 14 | S | Psocoptera | Geographic distance | 0.216 | 0.433 | 67.731 | <0.01 |
| 14 | S | Psocoptera | Null | 0 | 0.212 | 255.822 | <0.01 |
| 51 | S | Psocoptera | Temp | 0.552 | 0.691 | 0 | 0.61 |

| | | | | | | | |
|---|---|---|---|---|---|---|---|
| 51 | S | Psocoptera | Temp + Precip | 0.556 | 0.695 | 0.91 | 0.387 |
| 51 | S | Psocoptera | Precip | 0.622 | 0.804 | 10.825 | 0.003 |
| 51 | S | Psocoptera | Geographic distance | 0.226 | 0.451 | 68.774 | <0.01 |
| 51 | S | Psocoptera | Null | 0 | 0.22 | 270.135 | <0.01 |
| 80 | S | Psocoptera | Temp | 0.549 | 0.693 | 0 | 0.772 |
| 80 | S | Psocoptera | Temp + Precip | 0.549 | 0.693 | 2.712 | 0.199 |
| 80 | S | Psocoptera | Precip | 0.577 | 0.748 | 6.564 | 0.029 |
| 80 | S | Psocoptera | Geographic distance | 0.219 | 0.442 | 67.891 | <0.01 |
| 80 | S | Psocoptera | Null | 0 | 0.218 | 260.745 | <0.01 |
| 90 | S | Psocoptera | Temp | 0.532 | 0.665 | 0 | 0.779 |
| 90 | S | Psocoptera | Temp + Precip | 0.531 | 0.665 | 2.57 | 0.216 |
| 90 | S | Psocoptera | Precip | 0.498 | 0.639 | 10.012 | 0.005 |
| 90 | S | Psocoptera | Geographic distance | 0.222 | 0.431 | 65.328 | <0.01 |
| 90 | S | Psocoptera | Null | 0 | 0.203 | 257.625 | <0.01 |
| 92 | S | Psocoptera | Temp | 0.549 | 0.707 | 0 | 0.738 |
| 92 | S | Psocoptera | Temp + Precip | 0.548 | 0.705 | 2.69 | 0.192 |
| 92 | S | Psocoptera | Precip | 0.54 | 0.712 | 4.711 | 0.07 |
| 92 | S | Psocoptera | Geographic distance | 0.209 | 0.438 | 66.555 | <0.01 |
| 92 | S | Psocoptera | Null | 0 | 0.223 | 250.476 | <0.01 |

#### Supplementary Figures

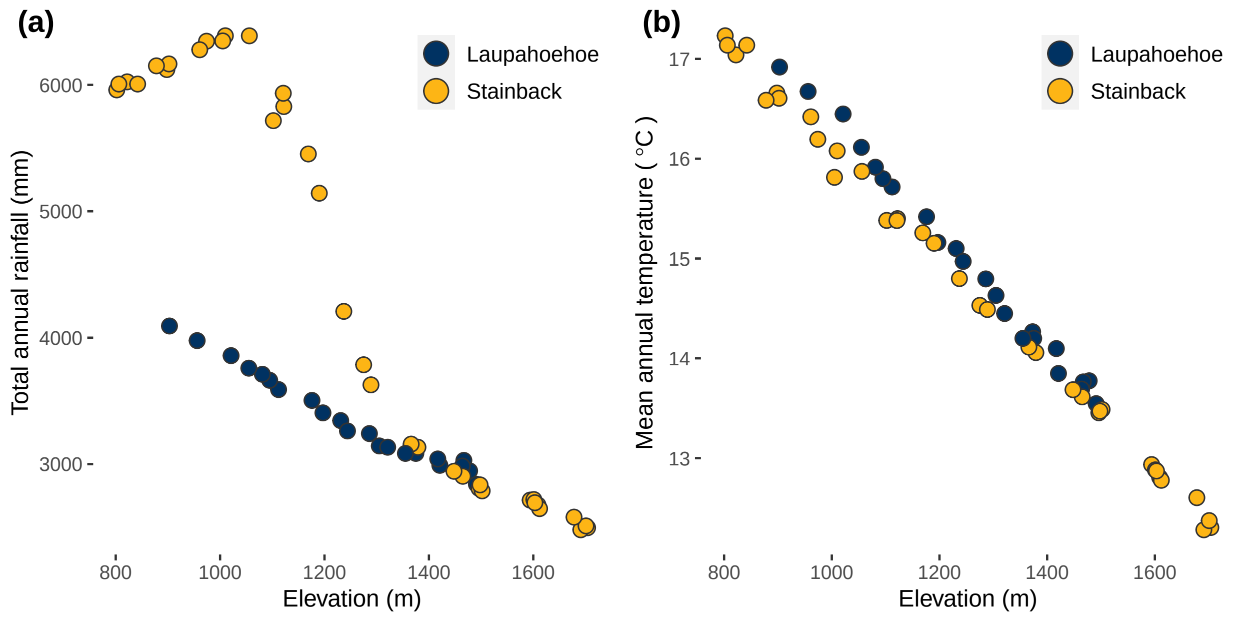


**Supplementary Figure 1: Precipitation and temperature with elevation along the two sampled transects along Mauna Kea and Mauna Loa.**

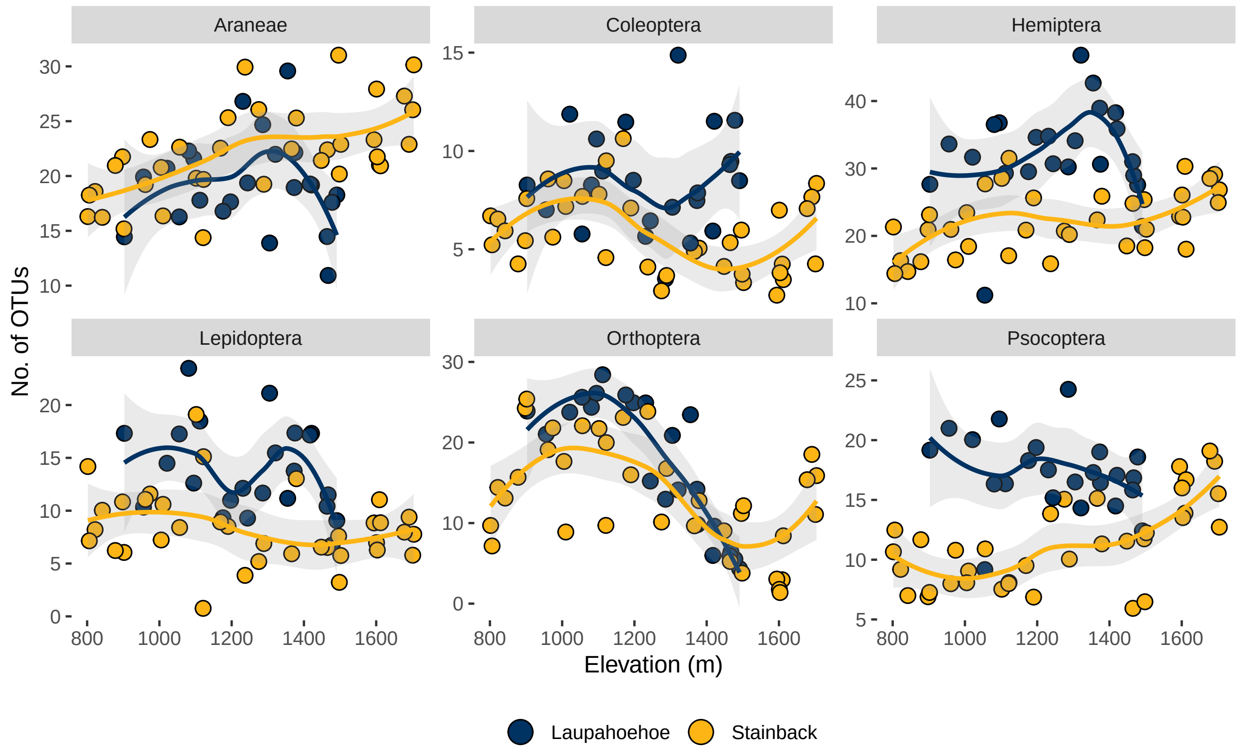


**Supplementary Figure 2**: **Taxonomic order-level OTU richness across two elevational transects on the volcanoes Mauna Kea (Laupāhoehoe) and Mauna Loa (Stainbeck) on the island of Hawai'i.** Points represent the average OTU richness for each site across rarefied datasets, whereas error bars represent the minimum and maximum OTU richness across rarefied datasets. Lines (mean) and shaded area (95% confident intervals) depict trends in OTU richness for each transect and were generated using a loess smoothing function on average OTU richness of sites in each transect. OTU richness of Coleoptera, Hemiptera, Lepidoptera and Psocoptera was significantly higher in Laupāhoehoe compared to Stainback (Welch's t-test, $p < 0.001$). However, the reverse is true for Araneae (Welch's t-test, $p = 0.02$). OTU richness of Orthoptera was not significantly different between transects (Welch's t-test, $p = 0.06$).

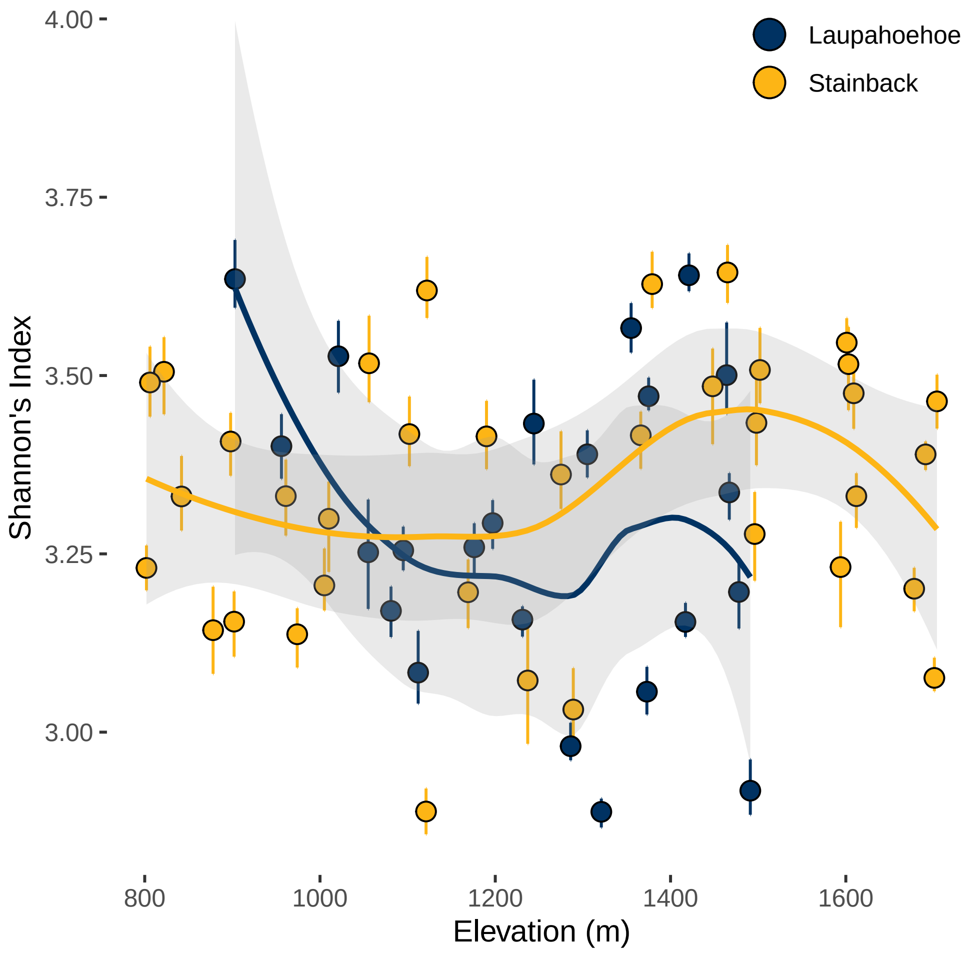


**Supplementary Figure 3: Shannon index of OTUs against elevation for both Laupāhoehoe (blue) and Stainback (yellow) transects.** Points represent the average Shannon index for each site across rarefied datasets, whereas error bars represent the minimum and maximum OTU richness across rarefied datasets. Lines (mean) and shaded area (95% confident intervals) depict trends in Shannon index for each transect and were generated using a loess smoothing function on average Shannon index of sites in each transect.

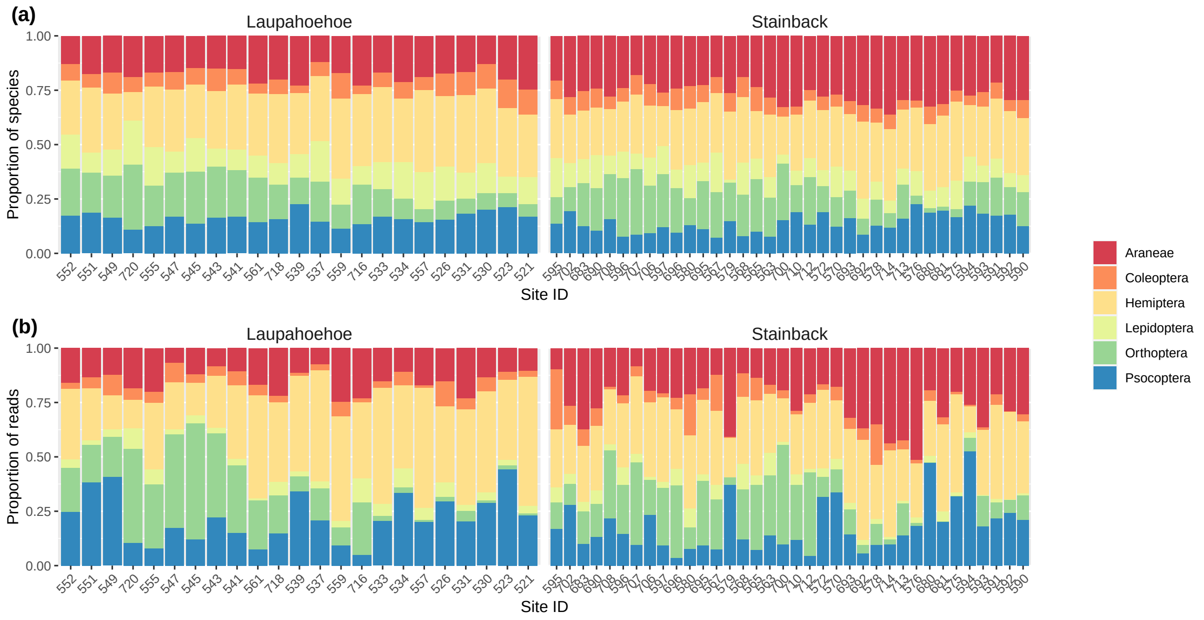


**Supplementary Figure 4: Proportion of species (a) and proportion of rarefied reads (b) in each taxonomic order at each site on Laupāhoehoe and Stainback transects.** Values are averages across 100 rarefied datasets. Sites are arranged in ascending order of elevation (lowest on left)

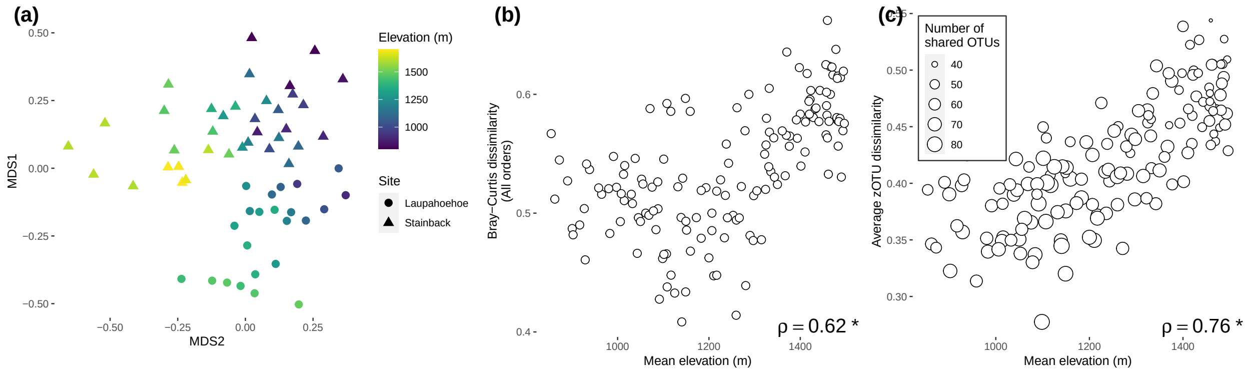


**Supplementary Figure 5**: **The effect of elevation on community compositional dissimilarity (unweighted Bray-Curtis dissimilarity) between sites on Laupāhoehoe vs. Stainback transects.** a) Non-metric dimensional scaling analysis of sampling localities across sites (triangles = Stainback, circles = Laupāhoehoe). NMDS stress value = 0.18. b) Pairwise dissimilarity between Laupāhoehoe and Stainback sampling localities against the mean elevation of localities compared. Only sampling sites between transects that were within 100 meters elevation of each other were compared. Arthropod communities are increasingly dissimilar between Laupāhoehoe and Stainback with higher elevation (Spearman's correlation, $\rho = 0.62$, $p < 0.05$ ). c) Average unweighted dissimilarity in zOTU composition of OTUs between sites on different transects increases with increasing elevation ($\rho = 0.76$, $p < 0.05$ ).

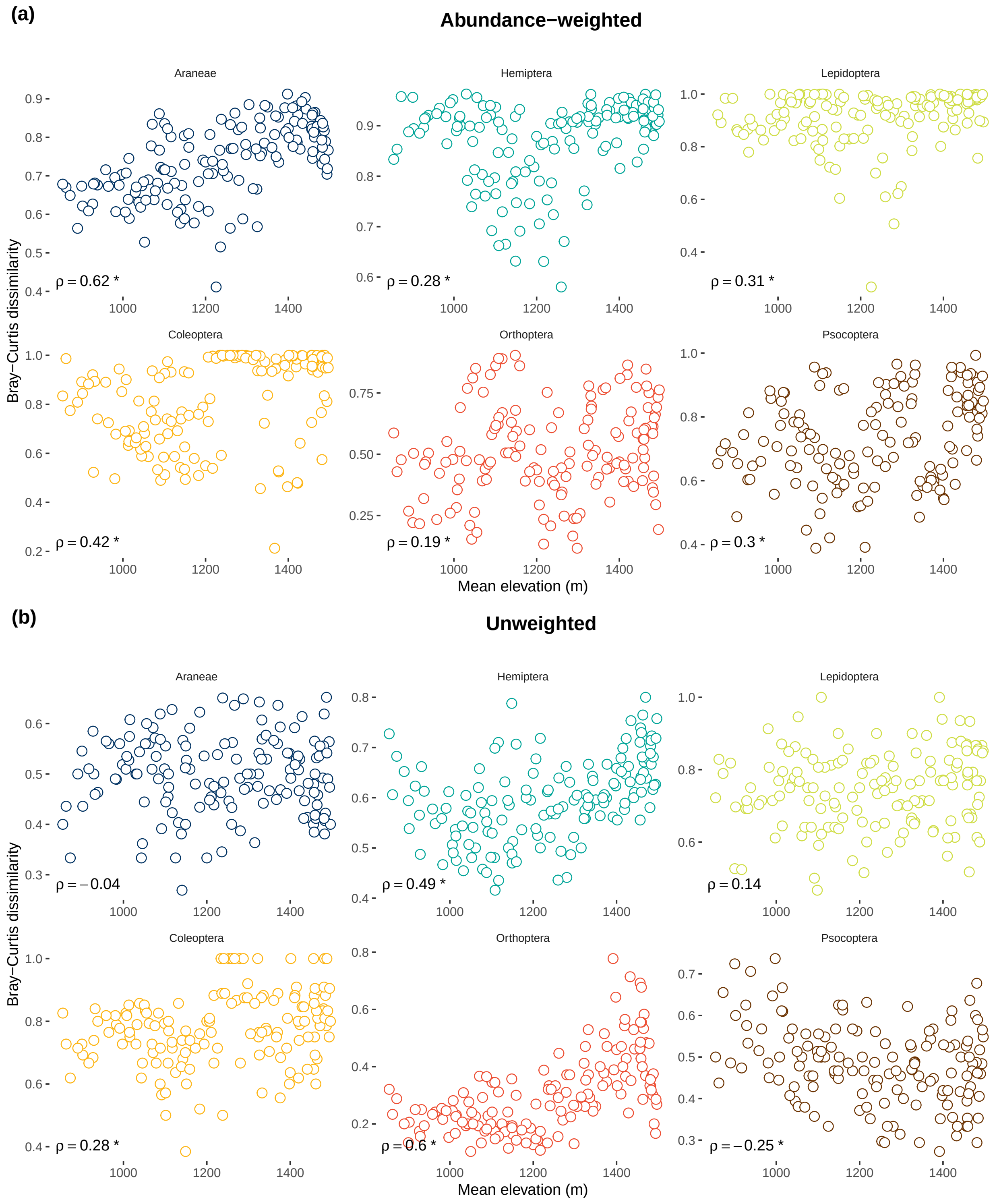


**Supplementary Figure 6**: **The effect of elevation on order-level OTU composition across both elevation gradients, based on (a) weighted and (b) unweighted Bray-Curtis dissimilarity.** Circle repreent pairwise unweighted dissimilarity values between Laupāhoehoe and Stainback sampling localities against the mean elevation of localities compared. Only sampling sites between transects that were within 100 meters elevation of each other were compared. Order-level assemblages are increasingly dissimilar between Laupāhoehoe and Stainback with higher elevation. $\rho$ values represent Spearman correlation between dissimilarity and elevation, * indicates that p-value $< 0.05$.

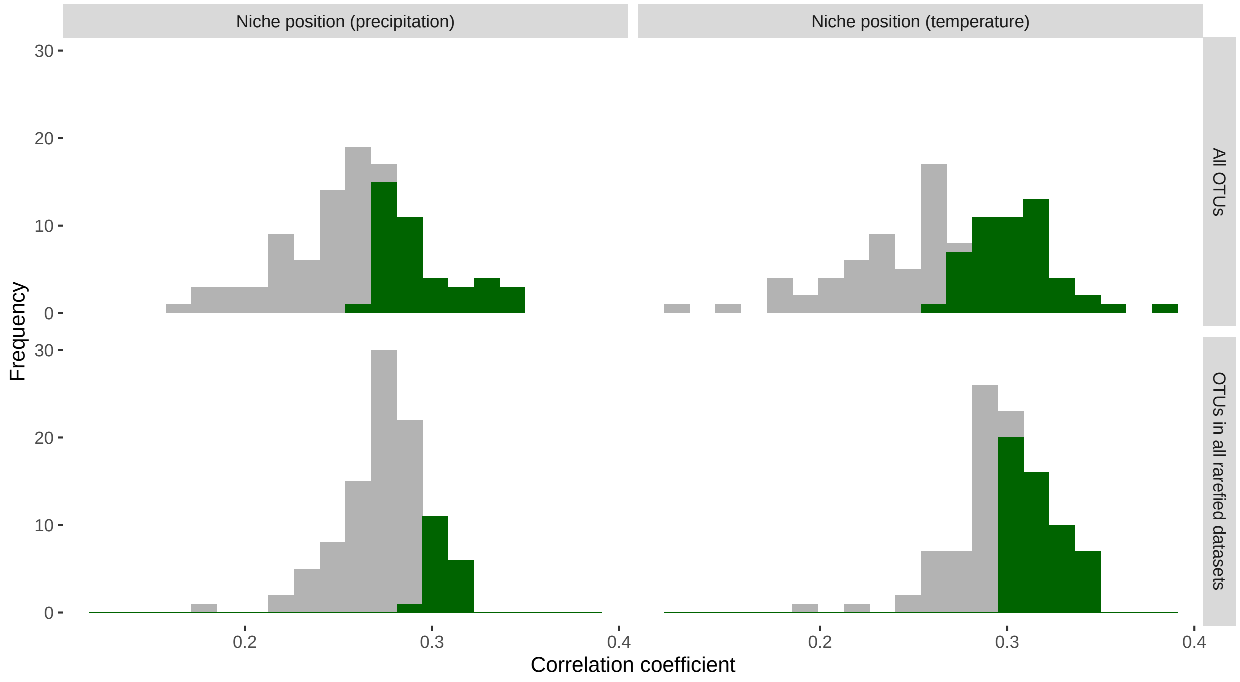


**Supplementary Figure 7**: **Correlation of climatic niches of arthropod OTUs between transects.** Histograms represent Spearman's correlation coefficients of climatic niche position across 100 rarefied datasets for all OTUs (top panels) and OTUs that occur in all rarefied datasets (bottom panels). Green bars represent the proportion of tests that were significant ($\alpha = 0.05$) based on permutation of values. Analysis was performed on OTUs that occurred in at least 10 sites on each transect.
